# MBOAT2 limits the amounts of PUFA in phosphatidylcholine of neonatal and dystrophic skeletal muscle and promotes muscle regeneration

**DOI:** 10.1101/2025.11.02.686003

**Authors:** William J. Valentine, Suzumi M. Tokuoka, Tomomi Hashidate-Yoshida, Keisuke Yanagida, Tomoyuki Suzuki, Natsuko F. Inagaki, Norio Motohashi, Kenta Nakano, Tadashi Okamura, Yoshihiro Kita, Hideo Shindou, Yoshitsugu Aoki, Takao Shimizu

## Abstract

Skeletal muscle regeneration is critically shaped by lipid remodeling, yet regulatory mechanisms involved remain unexplored. We identify MBOAT2, a lysophospholipid acyltransferase highly induced in activated satellite cells, as a key enzyme that limits the amounts of polyunsaturated fatty acids (PUFAs) in phosphatidylcholine (PC). Lipidomic profiling reveals a characteristic PC signature—high in monounsaturated fatty acids (MUFAs) and low in PUFAs—in dystrophic, injured, and neonatal muscle, paralleling elevated *Mboat2* expression. Given the susceptibility of PUFAs to oxidative damage and the emerging role of lipid peroxidation in driving muscle atrophy during denervation, disuse, and aging, MBOAT2-mediated enrichment of MUFA-PC may represent a protective and pro-regenerative lipid environment. Loss- and gain-of-function approaches confirm that MBOAT2 remodels PC and promotes myogenic repair. Our findings uncover a lipid remodeling circuit in muscle stem cells that buffers oxidative stress and highlight that MBOAT2 may improve regenerative capacity in muscular dystrophy, sarcopenia, and other muscle-wasting conditions.

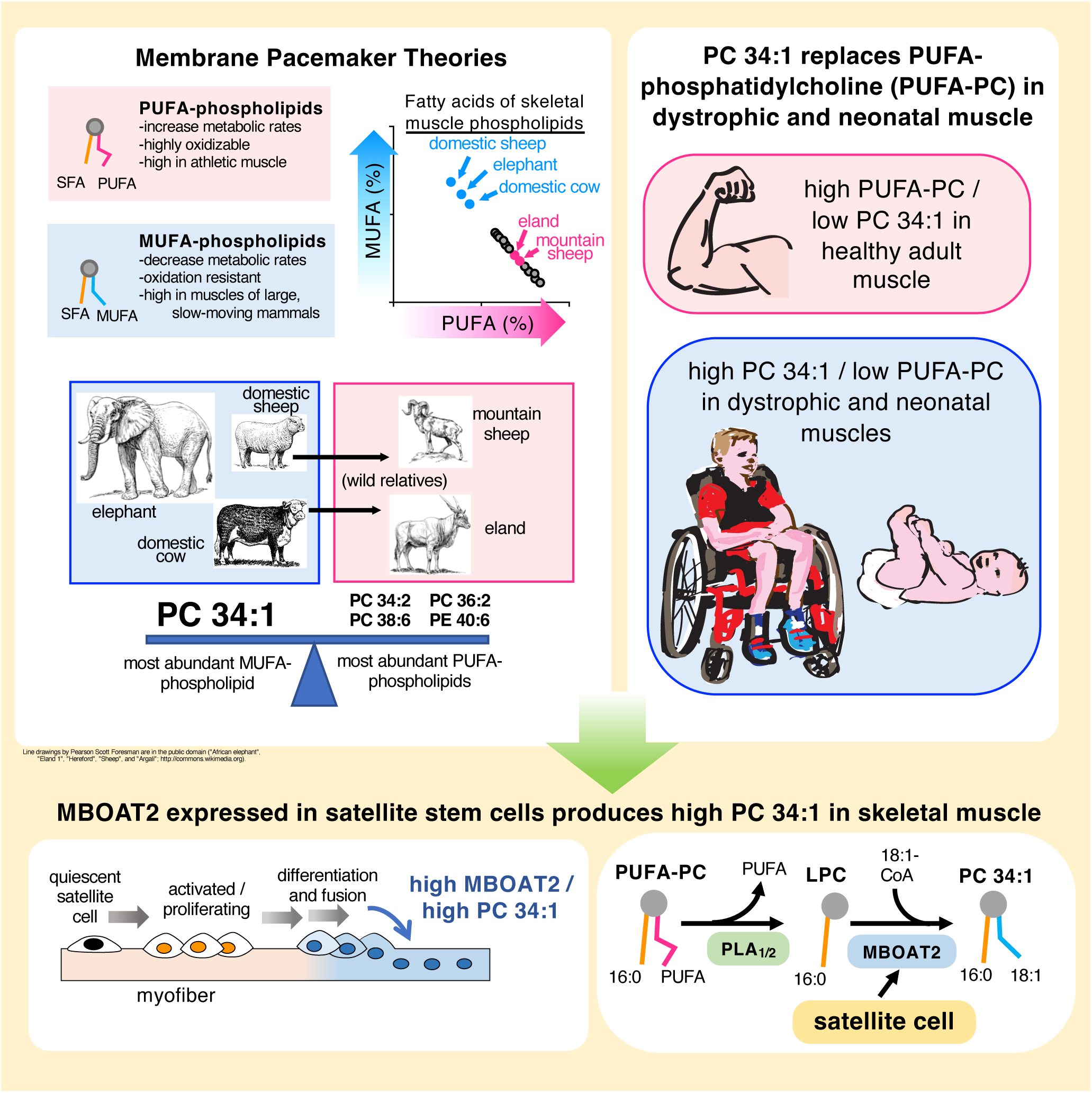

## INTRODUCTION

Phosphatidylcholine (PC) accounts for ∼half of all phospholipid in skeletal muscle (Figure 1A) and commonly contains a saturated fatty acid (SFA) at one acyl chain position and either monounsaturated fatty acid (MUFA) or polyunsaturated fatty acid (PUFA) at the other. Bis-allylic carbons of PUFAs possess nearly unrestricted rotational freedom [1] and are highly vulnerable to oxidation. In contrast, allylic carbons of MUFAs are structurally stable and resist oxidation (Figure 1B). PUFAs are main cellular targets to receive and propagate oxidative damage in lipid peroxidation chain reactions [2, 3], and MUFAs, when they replace PUFAs, can inhibit this process [4].

**Figure 1.**
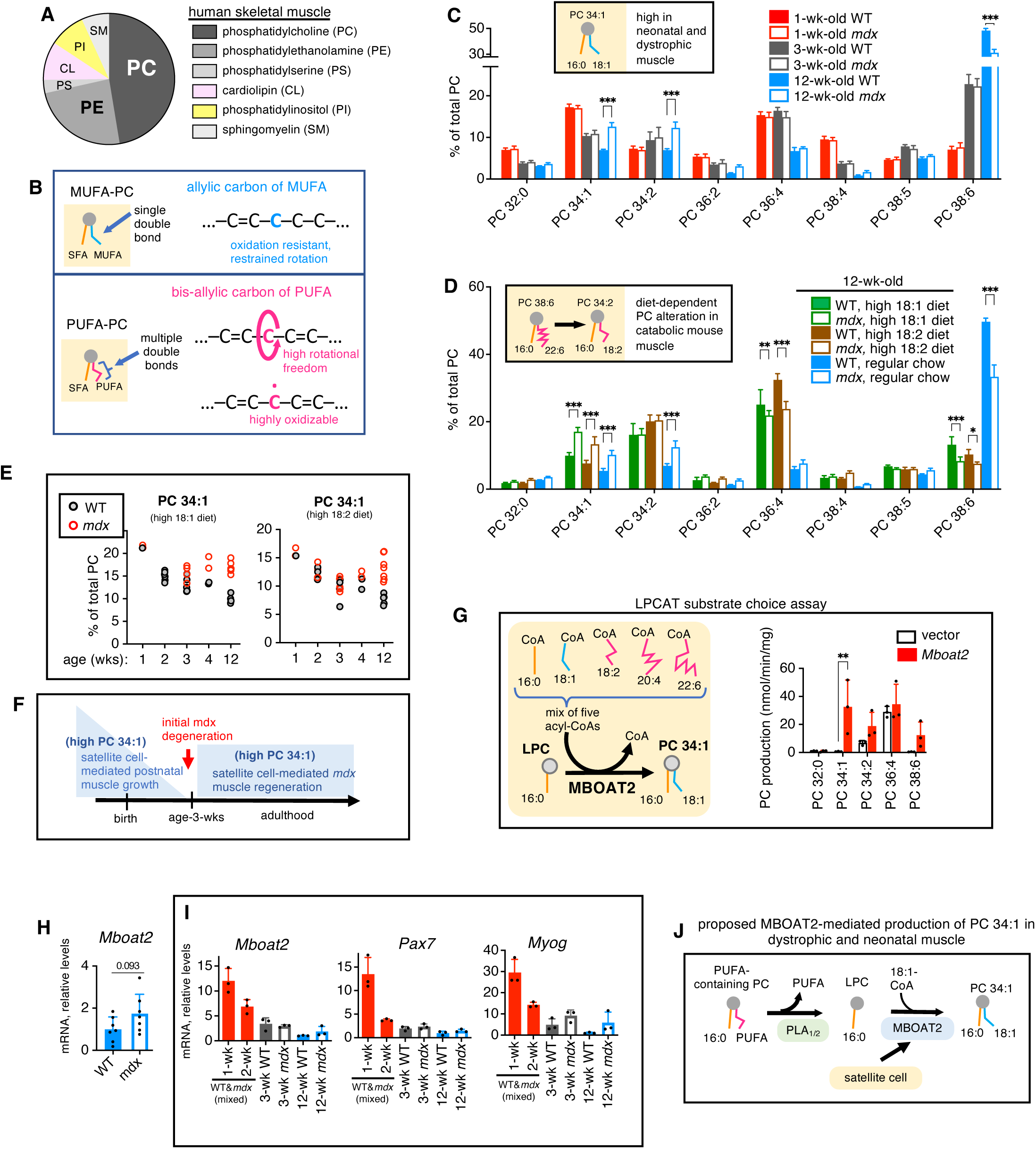
PC 34:1 and *Mboat2* expression levels in neonatal and dystrophic *mdx* mouse muscle. (A) Phospholipid compositions of human gastrocnemius muscles. Averages values of 10 males, 5- to 55-years-old. Plotted data represent values originally reported by Bruce [84]. (B) Schematic of typical MUFA-containing (upper panel) and PUFA-containing (lower panel) PC molecules. (C) PC compositions of gastrocnemius muscles of B10-WT and -*mdx* mice, ages 1-, 3-, or 12-weeks (n = 3–8 / group). Inset: schematic of high PC 34:1 in neonatal or dystrophic mouse muscle. Molecular species > 5% of total PC are plotted. (D) PC compositions of gastrocnemius muscles of 12-week-old WT and *mdx* mice raised on three different diets (n = 4–7 / group). For diet compositions, see Tables S3 and S4. Inset: schematic of non-specific, diet-dependent PC alterations in catabolic mouse muscle. Molecular species > 5% of total PC are plotted. (E) PC 34:1 levels in gastrocnemius muscles of B10-WT and -*mdx* mice, ages 1-, 2-, 3, 4- and 12-weeks, raised on two custom diets. Individual data points are plotted. Female tissues including unaffected carriers (counted as WT) were included in 1- to 4-week-old time points in order to increase sample number. For detailed gender and genotype information of each point, see Figure S1E. (F) Schematic of satellite cell activity coincides with PC 34:1 regulation in neonatal or dystrophic mouse muscle. (G) Biochemical substrate choice assay. Microsomes from vector- or *Mboat2*-transfected CHO-K1 cells were incubated with LPC 16:0 and a mixture of five different acyl-CoAs (n = 3 independent experiments). (H) *Mboat2* expression in gastrocnemius muscles of 12-week-old B10-WT and -*mdx* mice (n = 7 / group). (I) Expression levels of *Mboat2*, *Pax7*, and *Myogenin* in gastrocnemius muscles of 1-, 2-, 3-, and 12-week-old B10-WT and -*mdx* mice (n = 3 / group). 1- and 2-week-old samples are mix of genders (male and female) and genotypes (WT and *mdx*), which is not expected to affect the PC compositions at these very young ages. Includes data also plotted in panel H (12-week-old samples). (J) Schematic of proposed MBOAT2-mediated production of PC 34:1 in dystrophic or neonatal muscle. Error bars represent SD (C,D, G-I). Significance is based on t-test (H) or Bonferonni’s tests (C,D,G).

Membrane pacemaker theories of aging have noted phospholipid unsaturation is greater in smaller animals, which tend to have higher metabolic rates and shorter lifespans than larger animals, and propose that fatty acid compositions of membranes are key determinants of animal longevity based on the tendencies of PUFAs to both increase membrane-associated metabolic rates and propagate oxidative damage [5]. In comparative analyses of mammalian species, skeletal muscle phospholipid unsaturation negatively correlated with body mass and maximum lifespans [6, 7], MUFA and PUFA levels showed highly inverse relationships, and increased MUFA levels were associated with both increased body mass [6] and decreased (body-size-adjusted) maximum running speeds [8]. These observations suggest that balances of MUFA and PUFA in muscle may be under evolutionary pressures, with high MUFA levels associated with large, long-lived species providing resistance to lipid peroxidation at possible costs in locomotor capabilities [8, 9]. However, regulatory mechanisms that control balances of MUFA and PUFA in muscle, which greatly differ among mammalian species, have been unknown.

In mitochondria of multiple mammalian tissues, PC 34:1 (with 16:0_ 18:1 chain sets), was the most abundant MUFA-containing phospholipid, and proposed to substitute for PUFA-containing phospholipids in membranes that are resistant to oxidative damage [10]. Fourteen mammalian lysophospholipid acyltransferases (LPLATs) (Table S1) determine acyl chain compositions of phospholipids both during *de novo* synthesis and Lands’ cycle acyl chain remodeling [11, 12] (Figure S1A). PC 34:1 is a major phospholipid produced during *de novo* synthesis [13], and its high levels in certain tissues were thought to signify lack of Lands’ cycle remodeling. However, PC 34:1 levels are highly upregulated in dystrophic muscles of Duchenne Muscular Dystrophy (DMD) patients [14–17] as well as during fetal and neonatal human muscle during development [18], and we hypothesized that membrane-bound O-acyltransferase 2 (MBOAT2, also called LPLAT13), a Lands’ cycle LPLAT with biochemical selectivity to generate 18:1-containing PC including PC 34:1 [19], may generate high PC 34:1 in these conditions.

Muscle stem cells (satellite cells) mediate fetal and neonatal muscle growth and adult muscle regeneration. We found that MBOAT2 is robustly upregulated by activated satellite cells and generates high levels of PC 34:1 which limit the amounts of PUFA-containing PC in neonatal, dystrophic, and injured mouse muscle. MBOAT2 expression was also increased in dystrophic muscles of DMD patients, suggesting conserved functions in human muscle. PUFA peroxidation contributes to dystrophic muscle pathogenesis [20–24] and promotes muscle atrophy under conditions of muscle denervation, disuse, unloading, and aging [25–28]. Our study indicates MBOAT2-mediated production of PC 34:1 is a key regulator of balances of MUFA and PUFA in muscle and a promising therapeutic target to ameliorate muscle diseases and muscle aging.

## RESULTS

### High PC 34:1 levels occur in dystrophic and neonatal mouse muscle

DMD is caused by mutational loss of dystrophin, a major structural protein of myofiber cell membranes. *Mdx* mice have a natural loss-of-function mutation of dystrophin and are a research model for DMD [29]. Although *mdx* pathology is mild compared to DMD, specific increase of PC 34:1 occurs in dystrophic muscles of both DMD patients and *mdx* mice [30–34] (Table S2). We utilized *mdx* mice to uncover the biological origin of high PC 34:1 in dystrophic muscle. DMD is an X-linked disease; therefore, male mice were used except where noted. PC compositions were measured in gastrocnemius muscles of C57BL/10 strain (B10) -WT and -*mdx* mice at age-1-week, before disease onset; age-3-weeks, during initial *mdx* degeneration; and age-12-weeks, when *mdx* pathology is mild and stable. No major *mdx*-associated PC alterations were detected in 1- or 3-week-old muscles. However, PC 34:1, high PC 34:2, and low PC 38:6 were evident in 12-week-old *mdx* muscles (Figure 1C). PC 34:1 was the most abundant PC molecule in 1-week-old muscles regardless of *mdx* status. Thus, high PC 34:1 is a common feature of both neonatal and dystrophic mouse muscle (Figure 1C, inset).

Increased PC 34:1 occurs in both *mdx* and DMD dystrophic muscle [16, 17, 30–34]; however, the increased PC 34:2 detected in *mdx* muscle contrasts to decreased PC 34:2 reported in DMD muscle [17]. PC of mouse muscle is more polyunsaturated than human muscle and also influenced by diet (Figures S1B-S1D), and 18:2-containing PCs like PC 34:2 are reported to be increased in catabolic mouse muscle in order to offset loss of 22:6 (DHA)-containing PCs like PC 38:6 [35] (Figure 1D, inset). Our regular mouse chow (Clea CE-2) is high in DHA. We raised mice on custom diets low in DHA but rich in either 18:1 or 18:2. These diets were also used in our previous study [34], nutritional contents are listed in Tables S3 and S4. PC 34:1 was consistently increased in dystrophic *mdx* muscle regardless of diet, while PC 34:2 alterations were diet-dependent (Figure 1D).

PC 34:1 levels in gastrocnemius muscles of B10-WT and -*mdx* mice raised on the custom diets were closely monitored during postnatal development (Figure 1E). One-to four-week-old muscles included tissues of both sexes, as no gender-dependent effects were evident at these ages (Figure S1E). Satellite cells actively grow neonatal mouse muscle for the first two weeks of life but nearly cease their activity by age-3-weeks [36–40]. However, satellite cells are selectively reactivated in *mdx* muscle from age ∼ 4-weeks in *mdx* muscle as they mediate regenerative responses [40]. PC 34:1 levels mirrored these activities, (Figures 1E and 1F), suggesting satellite cell-mediated pathways may generate high PC 34:1 in neonatal and dystrophic muscles.

### MBOAT2 produces PC 34:1 and is upregulated in neonatal and dystrophic muscle

MBOAT2 has biochemical selectivity to incorporate 18:1 into PC [19] but has no known function in skeletal muscle. We performed substrate choice biochemical assays by incubating membrane fractions of vector - or *Mboat2-*transfected CHO-K1 cells in reactions containing LPC 16:0, which is abundant in muscle [41], and a mixture of five acyl-CoAs (Figure 1G). MBOAT2 had clear preference to utilize 18:1-CoA and produce PC 34:1, suggesting MBOAT2 may produce PC 34:1 in muscle.

We measured *Mboat2* expression in gastrocnemius muscles of B10-WT and -*mdx* mice. *Mboat2* expression tended to be increased in 12-week-old B10-*mdx* muscles compared to B10-WT (Figure 1H). *Mboat2* expression was highly elevated in 1-week-old muscles and steeply decreased by age-3-weeks regardless of *mdx* status, similar to expression patterns of satellite cell markers *Pax7* and myogenin (*Myog*) (Figure 1I). These results suggest MBOAT2 upregulated by activated satellite cells may generate high PC 34:1 in dystrophic and neonatal muscle (Figure 1J).

### Expression of LPLATs during myogenic differentiation of satellite cells

Satellite cells reside along mature myofibers in quiescent states and in response to injury proliferate and fully differentiate, fusing to new or preexisting myofibers (Figure 2A). Primary satellite cultures isolated from C57BL/6 (WT) mice were induced to form myotubes by incubation in low serum media, and LPLAT expression levels were monitored. In initial screening, *Mboat2* and *Agpat3* were the most upregulated and *Lpcat1* was the most rapidly downregulated among all LPLATs (Figure 2B). LPCAT1 has enzymatic selectivity to incorporate 16:0 into PC and produce saturated PCs, PC 30:0 and PC 32:0, as well as PC 32:1 [42–46]. AGPAT3 promotes production of DHA-containing PC and phosphatidylethanolamine (PE) [47–50] and may have roles in metabolic adaptations of muscle to exercise [51].

**Figure 2.**
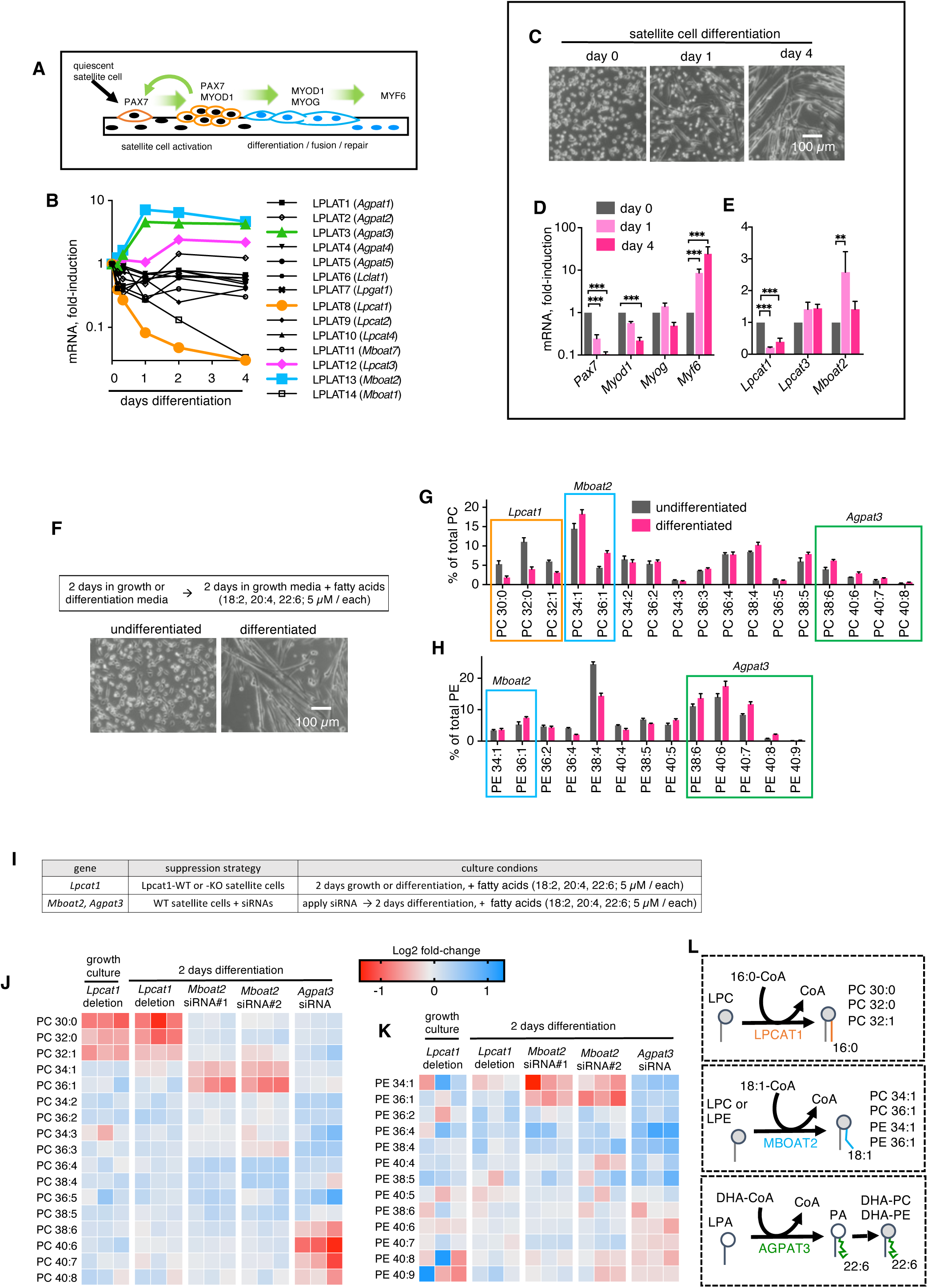
Expressions of LPLATs and phospholipid compositions of satellite cells. (A) Schematic of satellite cell-mediated muscle repair. (B) Screening of LPLAT1-14 expressions during primary satellite cell differentiation (n = 1 experiment). LPLATx nomenclature (where “x” is a number 1 to 14 [11]) is shown for each enzyme, with official gene names which are used throughout the present manuscript in parenthesis. (C-E) Primary satellite cells were differentiated for 4 days. Microscopic images of the cells (C) are shown. Mean expression levels of selected myogenic marker genes (D) and LPLATs (E) are plotted (n = 3 independent experiments). (F-H) Primary satellite cells either kept in growth media (undifferentiated) or differentiated for 2 days were cultured 2 additional days in growth media supplemented with fatty acids; representative images of the cells are shown (F). Compositions of PC (G) and PE (H) are plotted (n = 4 independent experiments). Candidate regulatory LPLATs are indicated for the boxed molecular species. (I-K) Effects of LPLAT gene suppression on PC and PE compositions of satellite cells. Schematic of gene suppression strategies is shown (I). Heatmaps summarize the effects of gene suppression on relative levels of individual molecular species of PC (J) and PE (K) (n = three independent experiments / condition). For complete data sets, see Figured S2E-S2I. (L) Schematics of activities of LPCAT1, MBOAT2, and AGPAT3 to regulate PC and PE in satellite cells. Error bars represent SEM (D, E, G, and H). Significance is based on Bonferonni’s tests (D-E).

We further examined expressions of *Lpcat1* and *Mboat2* as well *Lpcat3*, which incorporates 20:4 into PC and PE [52–55] (Table S5). During four-days differentiation to myotubes (Figure 2C), the satellite cell markers of quiescence (Pax7) and early activation (*Myod1*) were downregulated, the differentiation marker *Myog* was unchanged, and the late stage differentiation marker *Myf6* was increased (Figure 2D). During this time course, *Lpcat1* expression decreased, *Lpcat3* expression remained relatively constant, and *Mboat2* expression transiently increased (Figure 2E). Similar patterns of decreased *Lpcat1* expression, unchanged *Lpcat3* expression, and increased *Mboat2* expression occurred during differentiation of C2C12 myogenic cells (Figures S2A-S2C). These results suggest roles for LPCAT1 to function in proliferating myoblasts and MBOAT2 to function at later stages of myogenic differentiation.

### PC and PE compositions of satellite cells

PC and PE compositions were compared between undifferentiated satellite cells and those differentiated to form myotubes; cells were then cultured for two additional days in growth media supplemented with fatty acids (18:2, 20:4, and 22:6; 5 µM each) to provide abundant substrates for phospholipid synthesis [56] (Figures 2F). Known enzymatic products of LPCAT1 (PC 30:0, PC 32:0, and PC 32:1) tended to be decreased during differentiation, while those of MBOAT2 (PC 34:1, PC 36:1, PE 34:1 and PE 36:1) and AGPAT3 (PC 38:6, PC 40:6, PC 40:7, PC 40:8, PE 38:6, PE 40:6, PE 40:7, and PE 40:8) tended to be increased, suggesting LPCAT1, MBOAT2, and AGPAT3 may regulate PC and PE compositons of satellite cells (Figures 2G and 2H).

### LPLATs regulate PC and PE compositions of satellite cells

Genetic deletion and siRNA-mediated knock-down strategies were employed to examine the activities of LPCAT1, MBOAT2, and AGPAT3 to regulate PC and PE compositions of satellite cells (Figure 2I). In order to examine the activities of LPCAT1, satellite cells were isolated from the muscles of *Lpcat1*-WT and *Lpcat1*-global-gene-deficient mice (*Lpcat1*-KO mice) and PC and PE were analyzed in both in growth and differentiation conditions. In order to examine the activities of MBOAT2 and AGPAT3, WT satellite cells were incubated with two different *Mboat2*-specific siRNAs, an *Agpat3*-specific siRNA used previously [51], or non-targeting siControl. Each siRNA effectively blocked induction of its target (Figure S2D).

Each gene suppression strategy was employed in three independent experiments (Figures 2J, 2K, and S2E-S2I). *Agpat3*-targeting siRNAs tended to suppress levels of DHA-containing PCs and PEs (PC 38:6, PC 40:6, PC 40:7, PC 40:8, PE 40:6, PE 40:7, PE 40:8, and PE 40:9), consistent with our previous report [51]. *Lpcat1* deletion reduced PC 30:0, PC 32:0, and PC 32:1 under both growth and differentiation conditions, and *Mboat2*-targeting siRNAs suppressed levels of PC 34:1, PC 36:1, PE 34:1 and PE 36:1, revealing novel activities of LPCAT1 and MBOAT2 to regulate phospholipid compositions of satellite cells (Figure 2L).

### PC and PE compositions and LPLAT expression levels following muscle injury

In order to examine whether the LPLAT-mediated regulation of PC and PE detected in cultured satellite cells may also be detected in regenerating muscle tissues, tibialis anterior (TA) muscles of WT mice were injured by barium chloride injection, and contralateral muscles were controls. In this model, peak satellite cell proliferation occurs at ∼3 days post-injury (dpi) and satellite cell-mediated repair is completed 14 dpi [57]. Tissues were collected 3-21 dpi for lipidomic and gene expression analyses (Figure 3A).

**Figure 3.**
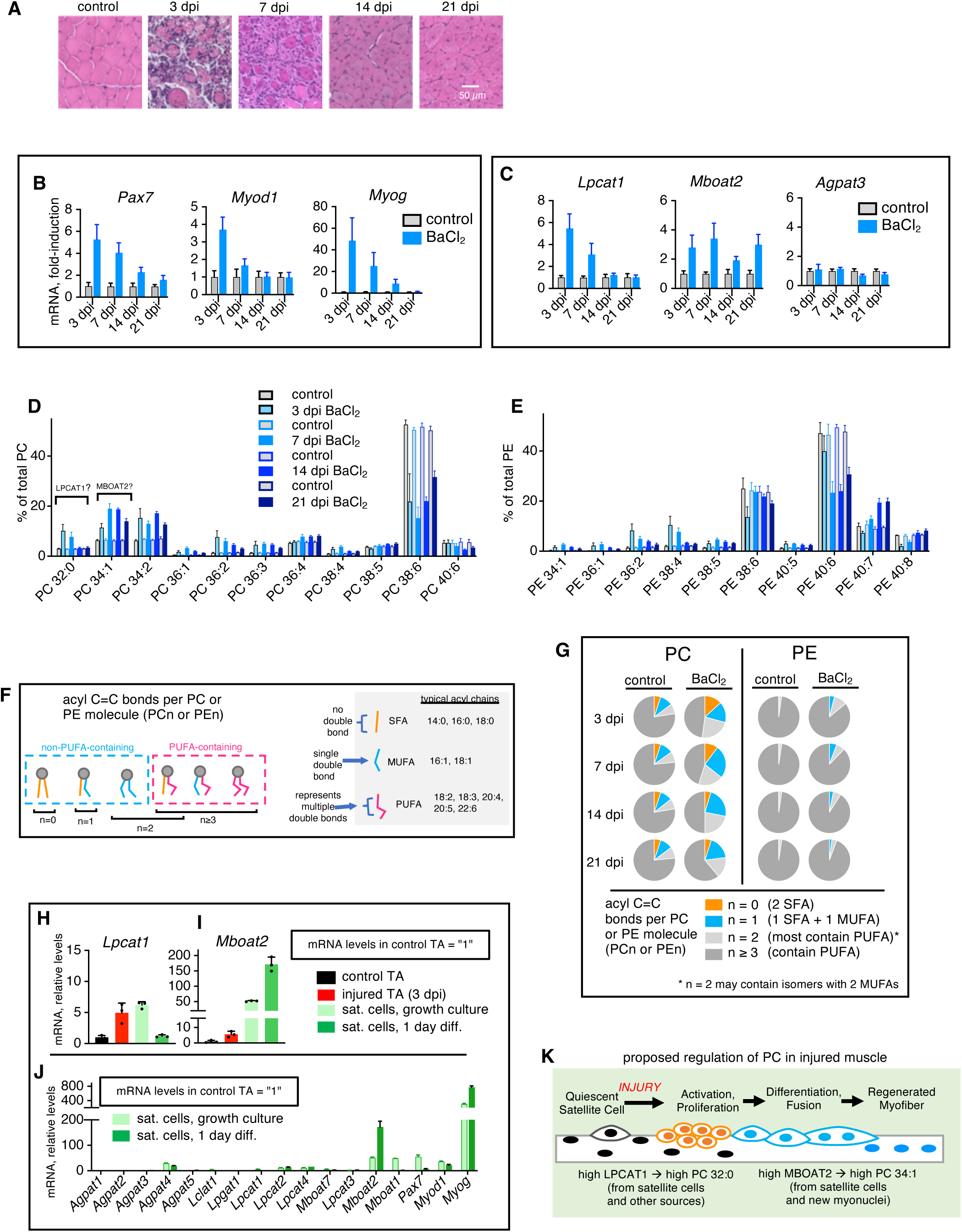
Gene expression and phospholipid compositions after muscle injury of WT mice. TA muscles were injected with barium chloride to induce muscle injury, and tissues were collected 3–21 days post-injury (dpi). Uninjected contralateral muscles were controls. (A) Representative images of H&E-stained muscle sections. (B-C) mRNA levels of myogenic marker genes (B) and selected LPLATs (C) (n = 5 / group). (D-E) Compositions of PC (D) and PE (E) were measured by LCMS (n = 5/group). Suggested regulation of PC 32:0, by LPCAT1, and PC 34:1, by MBOAT2, are indicated. (F) Schematic of PCn and PEn unsaturation indexes. “n” is the sum of C=C double-bonds in both acyl chains of each phospholipid. (G) PCn and PEn unsaturation indexes after muscle injury. Values were calculated from data sets plotted in panels D and E. (H-J) Comparative gene expressions in control and cardiotoxin-uninjured (3 dpi) TA muscles and primary satellite cells. LPLAT and myogenic marker gene expressions were normalized to *18S* as a reference gene in order to compare expression levels between muscle cells and tissues. Expression levels of each gene in control TA muscle were arbitrarily set to 1 following normalization to *18S*. Expression patterns of *Lpcat1* (H) and *Mboat2* (I) are plotted. Expressions in satellite cells relative to control muscle are plotted for all fourteen LPLATs and and three myogenic marker genes (J). RNAs from muscles (n = 3/group) were processed and analyzed in parallel with RNAs isolated from primary satellite cells. For complete analyses, see Figure S3A-S3D. (K) Schematic of proposed regulation of PC by LPCAT1 and MBOAT2 after muscle injury. Error bars represent SD (B-E, H-J).

Expression of *Pax7*, *Myod1*, and *Myog* was highest at 3 dpi followed by 7 dpi (Figures 3B) as was *Lpcat1* (Figure 3C, left), suggesting *Lpcat1* was predominately expressed in satellite cells and/or other cell types activated at these time points (i.e., immune cells, fibroblasts, endothelial cells). *Mboat2* expression was robustly increased throughout the time course (Figure 3C, center). *Agpat3* expression was relatively unaffected by muscle injury (Figure 3C, right).

PC and PE compositions were measured (Figures 3D and 3E). PC 32:0, was increased in injured muscle at 3 and 7 dpi, similar to *Lpcat1* expression, and PC 34:1, was robustly increased throughout the time course, similar to *Mboat2* expression, suggesting LPCAT1 generates increased PC 32:0 and MBOAT2 generates increased PC 34:1 in injured muscle.

PC or PE unsaturation can be described by acyl chain C=C double-bond number (PCn or PEn), the sum of double-bonds in both acyl chains (Figure 3F). Saturated PC (PCn = 0) was increased at 3 and 7 dpi and monounsaturated PC (PCn = 1) was increased throughout the time course, displacing PUFA-PC (Figure 3G, left). PE compositions were more polyunsaturated and less affected by injury than PC (Figure 3G, right).

Expression levels of LPLATs were compared between primary satellite cells and TA muscles; 18S was used as a stable reference gene for the comparisons (Figure S3A) [58]. TA muscles were injured by cardiotoxin injection, which induces similar regenerative responses as barium chloride[59], and control and injured tissues were collected 3 dpi. *Lpcat1* expression showed little enrichment in satellite cells compared to muscle tissues (Figure 3H), suggesting additonal cell types may also contribute to increased *Lpcat1* expression after muscle injury. *Mboat2* expression was remarkably enriched (>100-fold) in satellite cells compared to uninjured muscle (Figures 3I, 3J, S3B-S3D), suggesting satellite cells and their newly donated myonuclei are predominant sources of high *Mboat2* expression after muscle injury (Figure 3K).

### LPCAT1 contributes to increased PC 32:0 in injured muscle

To determine whether LPCAT1 generates PC 32:0 in injured muscle, TA muscles of *Lpcat1*-WT and *Lpcat1*-KO mice were injured by barium chloride injection and PC and PE compositions were measured 7 dpi (Figures 4A, 4B, S4A, and S4B). Injury-induced increase of PC 32:0 was partially suppressed in *Lpcat1*-KO muscle, while other major injury-associated PC alterations such as increased PC 34:1, increased PC 34:2, or decreased PC 38:6, were not diminished (Figure 4C). Less abundant enzymatic products of LPCAT1 in satellite cells, PC 30:0 and PC 32:1, were also selectively suppressed in injured *Lpcat1*-KO muscle (Figure S4B). These results indicate LPCAT1, possibly expressed in multiple cell types including satellite cells, contributes to increased PC 32:0 detected seven days after muscle injury.

**Figure 4.**
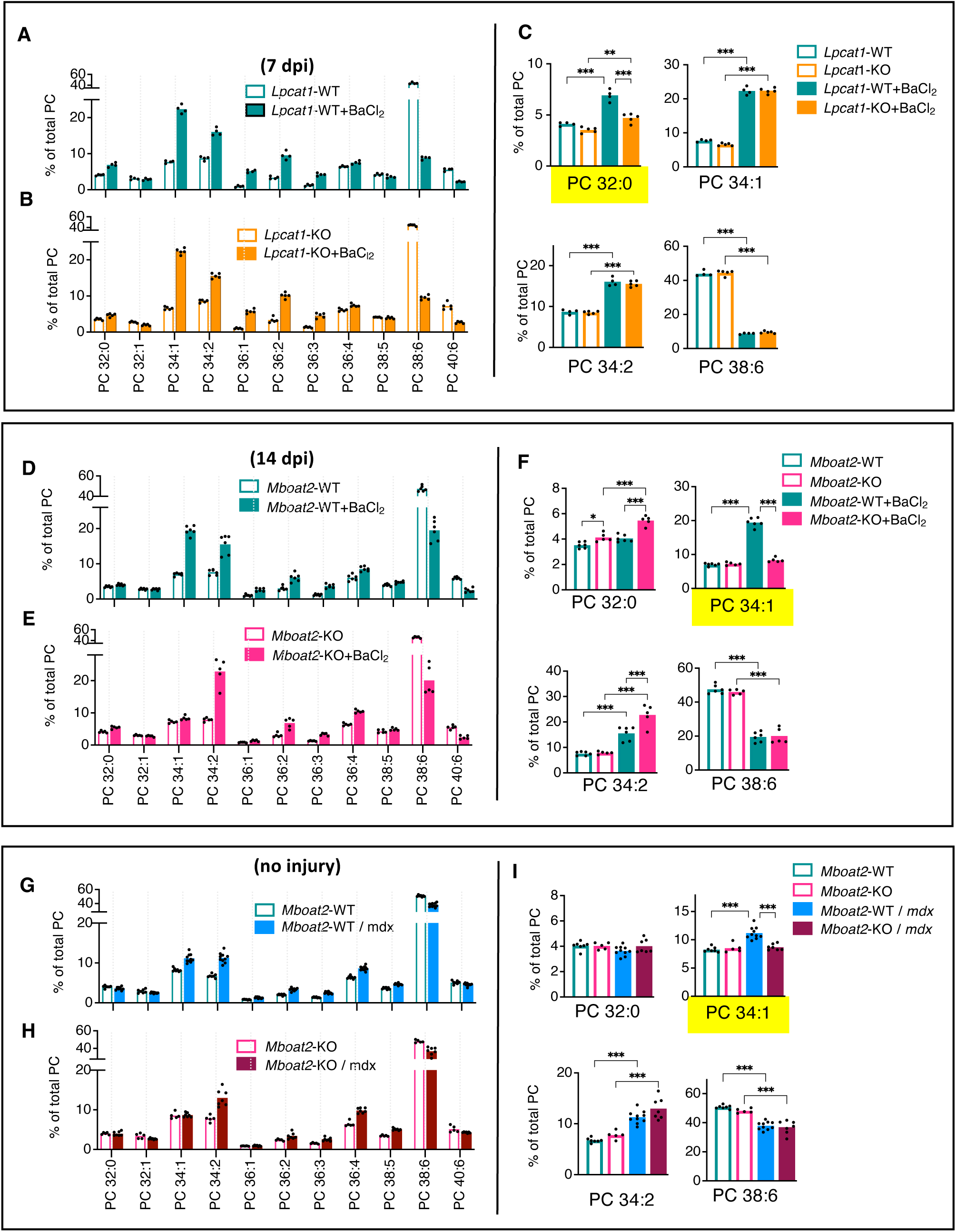
Comparative analyses of PC compositions in regenerating muscles of WT and *Lpcat1*- or *Mboat2*-KO mice. (A-D) TA muscles of 20-week-old *Mboat2*-WT and *Mboat2*-KO mice after injury by barium chloride injection (14 dpi), or on the dystrophic *mdx* background (non-injured). Muscle section were stained by H&E and representative images are shown (A). Minimum diameters of individual myofibers were measured in the muscle tissues, and myofiber diameter frequency distributions (B), average diameters (C), and variance coefficients (D) are plotted (n = 3–5 muscles / group; >200 fibers/muscle were measured). (E) Effects of *Mboat2* gene deficiency on PCn and PEn unsaturation indexes in control, injured, and dystrophic TA muscles. Values were calculated from the data sets plotted in Figure 4. (F) Fatty acid compositions of PC in quadricep muscles of DMD patients and controls. Plotted data represent values originally reported by Kunze et al. [15] and are thought to largely reflect replacement of PC 34:2 by PC 34:1 in DMD muscle [17]. (G) Estimated levels of non-PUFA and PUFA-containing PC and PE in mouse models of muscle regeneration and control and DMD human quadriceps. Estimations of PUFA-containing PC fractions in mouse muscle were calculated as the sum of PCn = 2 and and PCn ≥ 3 fractions plotted in panel E. Estimations of human muscle are based on data plotted in panel F and assume each PC or PE molecule contains at least one SFA or MUFA (PC with two PUFA chains are generally of low abundance [85, 86]). (H) *MBOAT2* expression levels in muscle biopsies from 10 DMD patients and 10 unaffected controls. Data were deposited in GEO data set GDS609 by Haslett et al. [61]. Error bars represent SD (B-D, F, H). Significance is based on t-tests (B-D, H).

### MBOAT2 generates increased PC 34:1 levels in injured muscle

To determine whether MBOAT2 produces high PC 34:1 detected in injured muscle, TA muscles of *Mboat2*-WT and *Mboat2*-global-gene-deficient (*Mboat2*-KO) mice were injured by barium chloride injection and PC and PE compositions were measured 14 dpi (Figures 4D, 4E, S4C, and S4D). Injury-induced increase of PC 34:1 was abolished in *Mboat2*-KO muscle, while other major injury-induced PC alterations including increased PC 32:0, increased PC 34:2, and decreased PC 38:6 were not diminished (Figure 4F). Less abundant enzymatic products of MBOAT2 in satellite cells, PC 36:1, PE 34:1, and PE 36:1, were also selectively suppressed in injured *Mboat2*-KO muscle (Figure S4D). These results indicate the high selectivity of MBOAT2 to increase PC 34:1 in injured muscle.

### MBOAT2 generates increased PC 34:1 levels in dystrophic *mdx* muscle

In order to determine whether MBOAT2 generates the increased PC 34:1 levels detected in dystrophic *mdx* muscle, *Mboat2*-KO mice and *mdx* mice (all on C57BL/6 backgrounds) were backcrossed. PC and PE compositions were measured in non-injured TA muscles of 20-week-old male Mboat2-WT, Mboat2-WT/*mdx*, *Mboat2*-KO, and *Mboat2*-KO/*mdx* mice (Figures 4G, 4H, and S4E). *Mdx*-associated increase of PC 34:1 was abolished in *Mboat2*-KO/*mdx* muscle while other major *mdx-*associated PC alterations including increased PC 34:2 and decreased PC 38:6 were not diminished (Figure 4I). Thus, similar MBOAT2-dependent increases of PC 34:1 were detected in both the injury and the dystrophic *mdx* models of muscle regeneration, but were of greater magnitude in the injury model.

### MBOAT2 is required for efficient regeneration after muscle injury

Histological examination of the TA muscles was performed to examine the effects of *Mboat2*-gene deficiency on muscle regeneration (Figures 5A-5D). Tissue architecture was similar in uninjured *Mboat2*-WT and *Mboat2*-KO muscles. However, *Mboat2*-KO muscles showed impaired regeneration two-weeks after barium chloride injection, including decreased myofiber diameters with greater variances, indicating an essential role of MBOAT2 for efficient regeneration following severe muscle injury. In contrast, *Mboat2* deletion did not strongly affect the mild dystrophic pathology of 20-week-old *mdx* mice (Figures 5A–5D).

**Figure 5.**
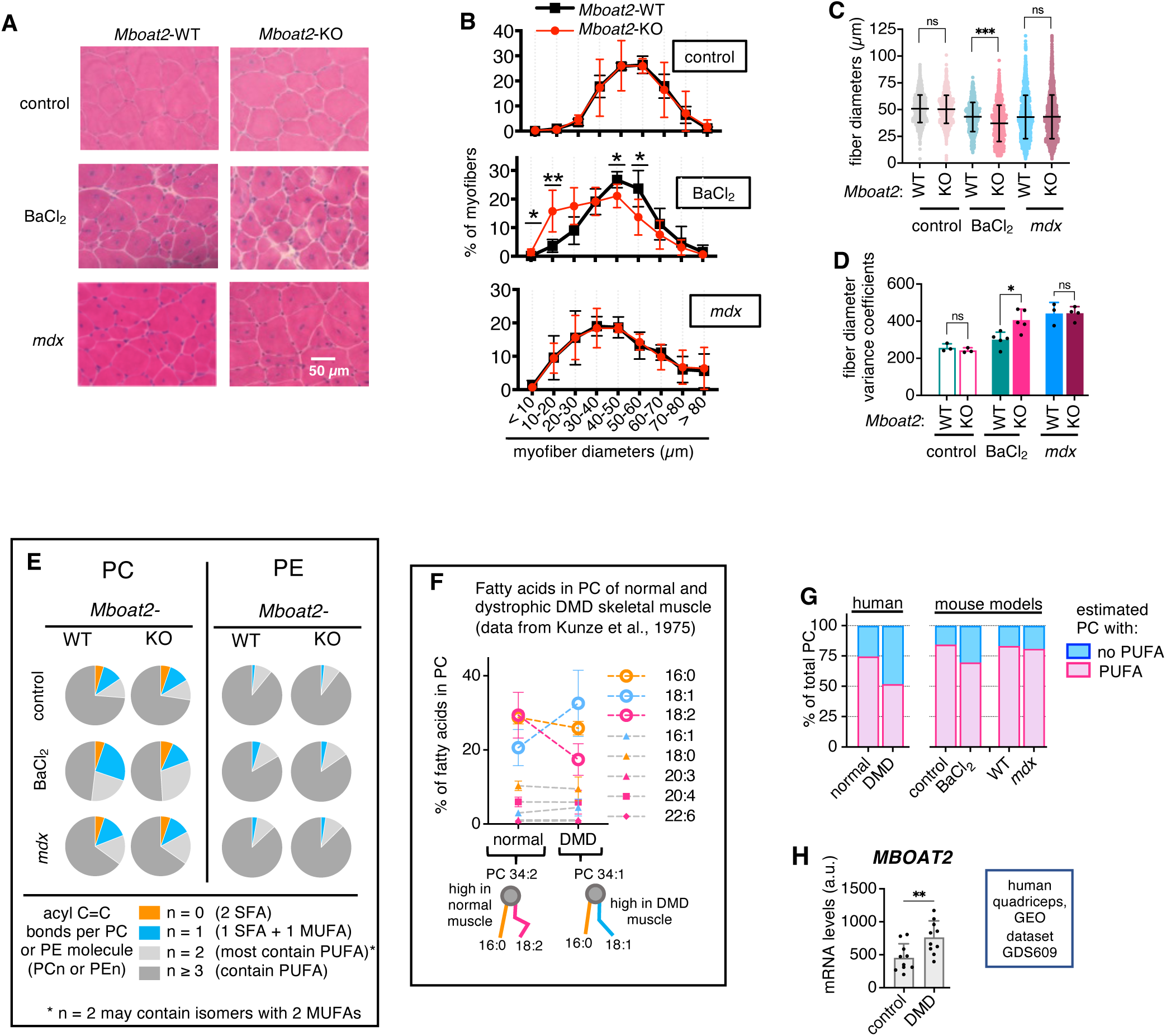
Evaluation of muscle pathology in injured or dystrophic muscles of *Mboat2*-WT and -KO mice. PC and PE compositions were measured in gastrocnemius muscles of male adult (1-year-old) and neonatal (1-week-old) *Mboat2*-WT and -KO mice. See Figures S5F and S5G for complete complete PC and PE analyses. (A-B) PC compositions of gastrocnemius muscles of male 1-year-old and 1-week-old *Mboat2*-WT (A) or *Mboat2*-KO (B) mice (n = 7-8 / group). (C) Selected data from panels A and B were replotted (n = 7-8 / group). Significance is based on Bonferonni’s tests. (D) Effects of *Mboat2* gene deficiency on PCn and PEn unsaturation indexes of 1-year-old and 1-week-old *Mboat2*-WT and -KO muscles. Values were calculated from the data plotted in Figures 6A, 6B, S5F, and S5G. (E) Fatty acid compositions of PC of human gastrocnemius muscles from gestation to middle age. Plotted data represent values originally reported by Bruce and Svennerholm [18]. (F) Estimated levels of non-PUFA and PUFA-containing PC based on data plotted in panel E. Estimations assume each PC or PE molecule contains at least one SFA or MUFA [85, 86]. (G) Schematic of proposed satellite cell- and MBOAT2-mediated regulation of PC 34:1 levels in neonatal muscle.

In *Mboat2*-WT mice, monounsaturated PC fractions (PCn = 1) were highly increased after muscle injury, but only modestly increased in the *mdx* dystrophic muscle model (Figure 5E, left panel). PE compositions were more unsaturated and less affected than PC in either model, suggesting MBOAT2 primarily regulates PC rather than PE in these models (Figure 5E, right panel).

Kunze et al. reported fatty acid compositions of PC in normal and DMD muscle [15] (Figure 5F). The increase of 18:1 and decrease of 18:2 is consistent with a large increase of PC 34:1 (and corresponding decrease of PC 34:2) that occurs in DMD muscle [16, 17]. Estimation of the magnitude of replacements of PUFA-containing PC in DMD muscle and the mouse muscle regeneration models we employed suggests the decrease of PUFA-containing PC in DMD muscle involves 24% of all PC molecules, while the decrease of PUFA-containing PC in *Mboat2*-WT mouse muscles involved 15% of all PC molecules in the barium chloride-induced injury mode, and just 2% of all PC molecules in the *mdx* muscular dystrophy model (Figure 5G).

That MBOAT2 was essential for efficient regeneration of severely damaged muscle following barium chloride injection but was non-essential in the context of *mdx* pathology suggests the magnitude of the MBOAT2-mediated PC alteration may be proportional to both the pathological burden and beneficial effects. Gene expression omnibus (GEO) [60] microarray data deposited by Haslett et al. [61] indicated MBOAT2 levels are increased in DMD muscle (Figure 5H), suggesting MBOAT2 may generate high PC 34:1 in DMD muscle by similar mechanisms as in dystrophic *mdx* or injured mouse muscle. The PC alteration in DMD muscle was quantitatively greater than either mouse model (Figure 5G), suggesting this alteration may be highly supportive of regenerative responses even if insufficient to overcome the severe DMD pathology caused by loss of dystrophin.

### MBOAT2 regulates high PC 34:1 levels in neonatal muscle

Satellite cells mediate both neonatal myogenesis and adult muscle regeneration [62]. To determine whether MBOAT2 mediates high PC 34:1 levels during neonatal muscle growth, we analyzed gastrocnemius muscles of adult (1-year-old) and neonatal (1-week-old) *Mboat2*-WT and *Mboat2*-KO mice. *Mboat2* expression was highly elevated in 1-week-old compared to 1-year-old muscle of *Mboat2*-WT mice (Figure S5A). *Mboat2*-KO mice had normal body weights at ages-3- and 4-weeks and normal expression patterns of *Pax7*, *Myod1*, or *Myog* in 1-week-old muscles, suggesting normal muscle growth (Figures S5B-S5E).

PC and PE compositions were measured (Figures 6A, 6B, S5F, and S5G). High PC 34:1 in 1-week-old *Mboat2*-WT muscle was specifically suppressed in 1-week-old *Mboat2*-KO muscle, while other PC characteristics of 1-week-old muscle such as increased PC 32:0, increased PC 36:4, and decreased PC 38:6 were not diminished (Figures 6C). Less abundant enzymatic products of MBOAT2 in satellite cells, PC 36:1, PE 34:1, and PE 36:1, also tended to be suppressed in 1-week-old *Mboat2*-KO muscles (Figure S5G). These results indicate the high selectivity of MBOAT2 to increase PC 34:1 in neonatal muscle.

**Figure 6.**
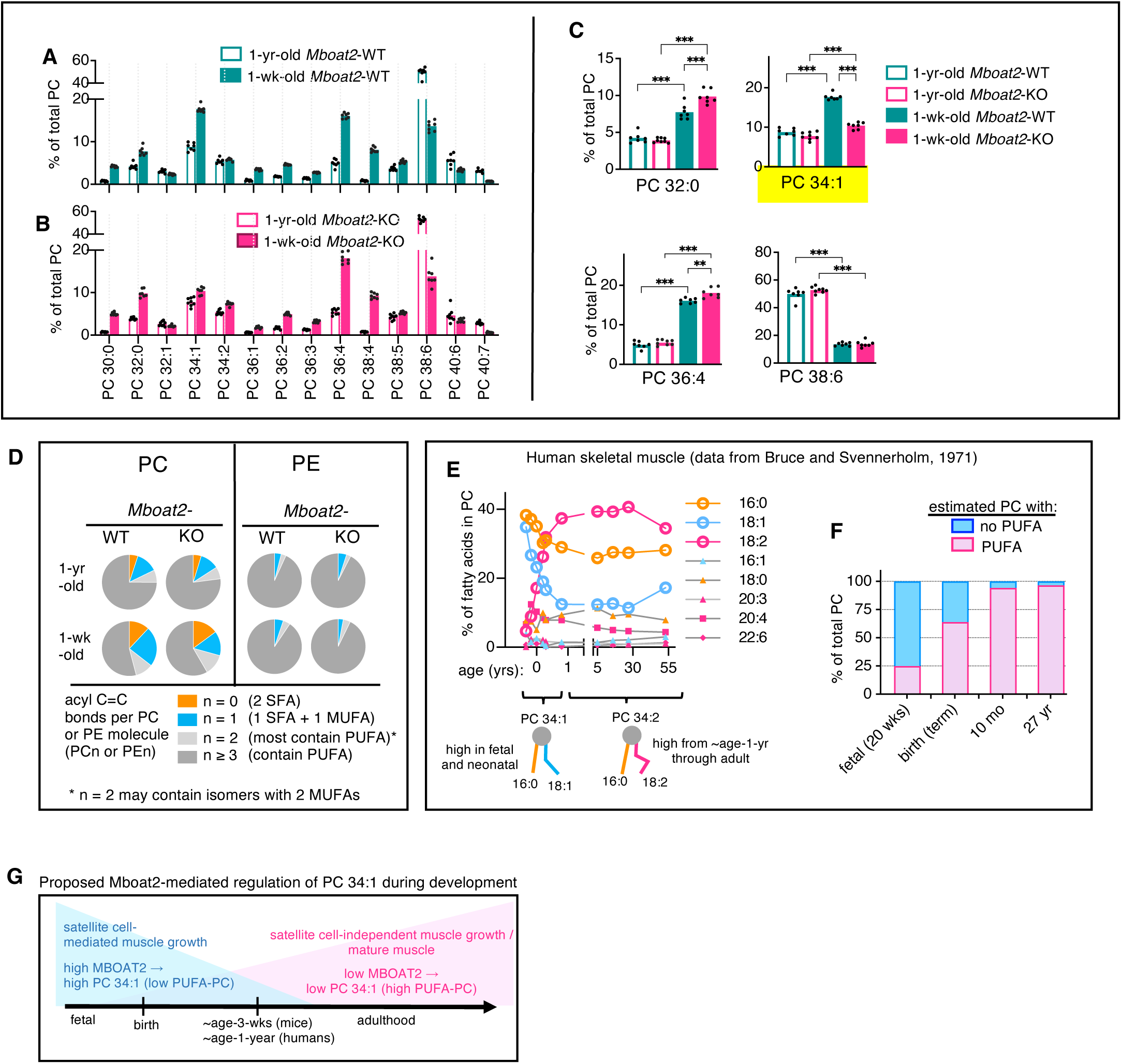
Effects of MBOAT2 on PC compositions of neonatal muscle. (A-C) TA muscles of *Lpcat1*-WT and -KO mice were injured by barium chloride injection and tissues were collected 7 dpi. Uninjected contralateral muscles were controls. PC compositions of uninjured and injured muscles were compared in tissues of *Lpcat1*-WT (A) or *Lpcat1*-KO (B) mice. Selected data from A and B were replotted (C). (D-F) TA muscles of *Mboat2*-WT and -KO mice were injected with barium chloride to induce muscle injury and tissues were collected 14 dpi. Uninjected contralateral muscles were controls. PC compositions of uninjured and injured muscles were compared in tissues of *Mboat2*-WT (D) or *Mboat2*-KO (E) mice. Selected data from D and E were replotted (F). (G-I) *Mdx* mice were crossed with *Mboat2*-KO mice, and PC compositions were measured in TA muscles of 20-week-old *Mboat2*-WT and *Mboat2*-KO mice, with or without the mdx mutation. PC compositions were compared in tissues of *Mboat2*-WT (G) or *Mboat2*-KO (H) mice. Selected data from G and H were replotted (I). Significance is based on Bonferroni’s tests (C, F, and I)(n = 4-10 / group). For complete comparative analyses of PC and PE between WT and KO, see Figure S4.

Both saturated PC (PCn = 0) and monounsaturated PC (PCn = 1) fractions were highly increased in 1-week-old compared to 1-year-old *Mboat2*-WT muscle; however, increased PCn = 1 was highly suppressed in 1-week-old *Mboat2*-KO muscle (Figure 6D, left). PE indexes were more polyunsaturated than PC and less affected by mouse’s age or *Mboat2*-deletion (Figure 6D, right). Bruce and Svennerholm [18] reported that high levels of 18:1 in PC of human fetal muscle are progressively and dramatically replaced by 18:2 chains by the first year of life, suggesting that high amounts of PC 34:1 are replaced by PC 34:2 during this period, dramatically increasing PUFA-PC (Figure 6E-F). Satellite cells have a critical role in postnatal muscle growth [62]. This activity is extremely high at birth but sharply declines during the first 3-weeks of life in mouse and first 1-2 years of life in humans [63–65], suggesting a broadly conserved function for MBOAT2 to generate high PC 34:1 during satellite cell-mediated stages of developmental muscle growth (Figure 6G).

### MBOAT2 generates high levels of PC 34:1 in differentiating satellite cells and myotubes

PC compositions of primary satellite cells isolated from WT and *Mboat2*-KO mice were measured in order to determine whether MBOAT2 contributes to high monounsaturated PC levels within newly formed myotubes. *Mboat2*-deficient satellite cells showed major deficits in PC 34:1 and PC 36:1 after four or six days of differentiation to myotubes (Figures 7A–7D), resulting in highly reduced monounsaturated PC (PCn = 1) levels in the *Mboat2*-KO myotubes (Figure 7E).

**Figure 7.**
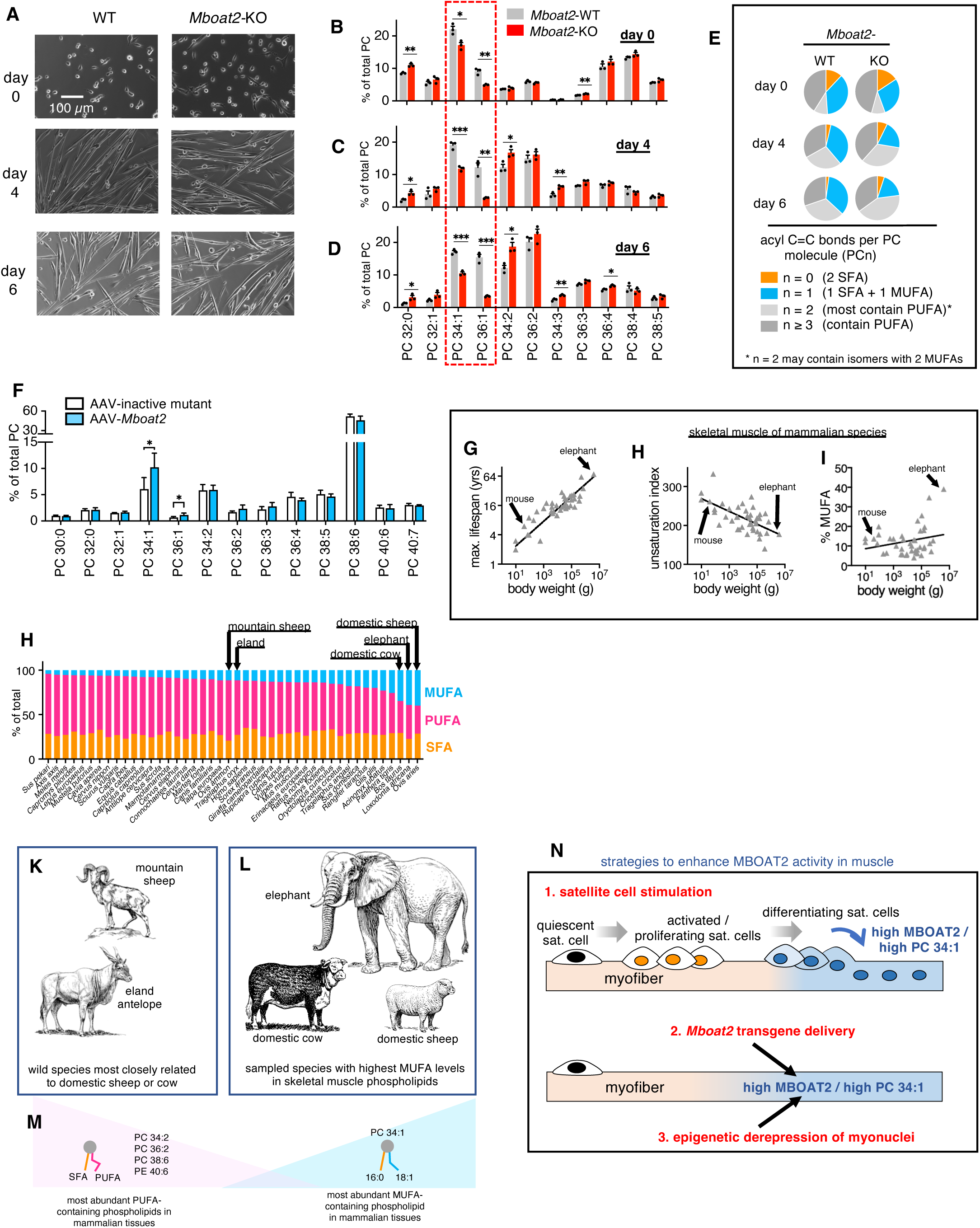
Effects of MBOAT2 on PC compositions of primary myotubes and adult mouse muscle. (A-E) Phospholipid compositions of primary satellite cells isolated by FACS from *Mboat2*-WT and *Mboat2*-KO muscle tissues. Represented images of the undifferentiated satellite cells (day 0) and myotubes at day 4 and day 6 of differentiation are shown (A). PC compositions of the undifferentiated satellite cells (day 0) (B) and differentiated myotubes after 4 (C) or 6 (D) days of differentiation. PCn unsaturation indexes (E) were calculated from the same data plotted in panel B-D. n=3 independent experiments. (F) Effects of AAV-*Mboat2* expression particles on PC compositions of mouse muscle. Gastrocnemius muscles of 5- to 8-week-old female B10-WT mice were injected with 5.7 x 10^10^ viral genomes (vg) of either AAV-inactive mutant *Mboat2* or AAV-*Mboat2* (n = 6/ group). Each injections also contained AAV-*eGFP* (1.4 x 10^10^ vg). See also Figure S6. (G-I) Correlative relationships of body weights of 42 mammalian specie to maximum lifespans (G), unsaturation index of skeletal muscle phospholipids (H), or amounts of MUFA as percentage of total fatty acids in skeletal muscle phospholipids (I). Plots were generated using data were first reported by Valencak et. al [82]. Unsaturation index is calculated as average number of C=C double-bonds per fatty acid multiplied times 100. Arrows indicate data for mouse and elephant as representative low and high body-weight species. (J) SFA, MUFA, and PUFA abundances in phospholipids of skeletal muscle of forty-four mammalian species, based on combined values originally reported in Valencak et al. [82] and Ruf and Valencak [8]. Samples were from hindlimb muscles, primarily musculus vastus, of adult specimens. For very small species, additional hindlimb muscle tissues were collected. Arrows indicate three species with highest MUFA amounts, African elephant, domestic cow, and domestic sheep, as well as the closest sampled wild relatives of the domestic cow, which is the eland antelope, or of the domestic sheep, which is the mountain sheep. (K-L) Line drawings of the selected mammalian species from panel J with highest MUFA levels in muscle (L) or their closest wild relatives included in the analyses (K). Line drawings by Pearson Scott Foresman are in the public domain (“African elephant”, “Eland 1”, “Hereford”, “Sheep”, and “Argali”; http://commons.wikimedia.org). (M) Schematic indicates most abundant PUFA-containing phospholipids (PC 34:2, PC 36:2, PC 38:6, and PE 40:6) and MUFA-containing phospholipid (PC 34:1) in multiple mammalian tissues, which are expected to significantly impact the MUFA and PUFA levels plotted in H [10]. (N) Schematic of three strategies to enhance MBOAT2 expression in adult muscle including (1) satellite cell stimulation as a natural method, and potential therapeutic strategies of (2) transgene delivery or (3) epigenetic derepression. Error bars represent SEM (B-D) or SD (F). Significance is based on t-tests (B-D and F).

### AAV-*Mboat2* increases PC 34:1 in healthy mouse muscle

*W*e tested whether ectopic AAV-*Mboat2* expression in uninjured adult mouse muscle may increase PC 34:1. Initial trials indicated injecting 5.7 x 10^10^ viral genomes (vg) of AAV-*Mboat2* induced robust *Mboat2* mRNA expression accompanied by elevated PC 34:1 levels (Figures S6A-S6D). This dose of AAV-*Mboat2* particles was injected into gastrocnemius muscles of adult female B10-WT mice, and AAV-inactive mutant *Mboat2* was injected into contralateral muscles as control. AAV-*Mboat2* particles induced pronounced increase of PC 34:1 (Figure 7F).

## DISCUSSION

The present study elucidates a novel pathway whereby activated satellite cells express LPLATs which limits PUFA in PC of neonatal or regenerating muscle. LPCAT1 was upregulated in proliferating satellite cells and produced PC 32:0. MBOAT2 was highly upregulated in differentiating satellite cells and generated high levels of PC 34:1. *Mboat2*-deficient mice selectively lacked increased PC 34:1 in conditions associated with high satellite cell activity including dystrophic, injured, and neonatal muscle, and showed impaired regeneration following muscle injury. Microarray data indicated MBOAT2 is highly elevated in DMD muscle, implicating MBOAT2 as the likely source of high PC 34:1 in DMD muscle. The disease course of DMD is prolonged, severe, and progressive [66], and decline of satellite cell function drives the progressive muscle wasting [67–70]. We propose that MBOAT2-generated high PC 34:1 may greatly support regenerative responses in early stages of DMD and interventions aimed at enhancing this pathway might improve the DMD disease course.

We identified MBOAT2 as the major source of high PC 34:1 in neonatal mouse muscle. The biological mechanisms that trigger onset of DMD and *mdx* dystrophic pathology are unknown. It is possible that MBOAT2-generated high PC 34:1 in neonatal muscle protects against dystrophic muscle disease onset, which is usually not detected before age 2-3 years in DMD patients and ∼age-3-weeks in *mdx* mice. Studies of patients in pediatric intensive care units suggested infants less than 1- or 1.5-years-old were resistant to muscle loss [71, 72], and whether neonatal muscle is resistant to muscle atrophy is an important area for future research.

Satellite cells have a major role in mediating the postnatal growth of mammalian skeletal muscle, with satellite cell density in the rapidly growing muscle generally dropping precipitously as a prelude to cessation of satellite cell activities and participation in muscle growth. Independently conducted genome-wide screens have identified MBOAT2 as among the most steeply downregulated genes during postnatal muscle growth of both domestic pigs [73] and cattle [74], suggesting MBOAT2-generated high PC 34:1 in neonatal muscle, reflective of satellite cell activity, may be widely conserved among mammals. A recent study indicated that MBOAT2 regulated the phospholipid compositions of neural stem cells and thereby promoted their abilities to activate [75], suggesting that MBOAT2-generated high PC 34:1 have a role to also promote activation of satellite cells. It has long been wondered why satellite cells stop their activity during later stages of postnatal muscle growth, and what are the biochemical cues which support their activation during earlier stages of fetal and postnatal muscle growth [76]. We speculate MBOAT2-generated MUFA-PC might provide one such cue, possibly influencing muscle sizes by affecting the timing or rate at which satellite cells cease their activity to participate in postnatal muscle growth.

Aging is a root element of all degenerative diseases, and longevity is a highly evolvable and positively selected evolutionary trait [77]. Membrane pacemaker theories of of aging propose that increased phospholipid unsaturation increases membrane associated metabolic rates as well as cellular susceptibility to oxidative damage, and by doing so increases the biological-rate-of living and decreases longevity. Body mass is an overarching physical trait that impacts most other physical parameters, and body mass of mammalian species shows strong positive correlations with maximum lifespans (Figure 7G) [78, 79]. Skeletal muscle comprises ∼40% of mammalian body mass [80], and body mass of mammalian species shows negative correlations with unsaturation of muscle phospholid and positive correlations with MUFA levels in muscle phospholipid (Figures 7H and 7I) [6]. However, major limitations of membrane pacemaker theories include that almost nothing is known of how acyl chain unsaturation of muscle phospholipids, which varies greatly among mammalian species, is regulated [81].

Analyses of phospholipids in skeletal muscle of forty-four mammalian species reported by Ruf et al. [8] and Valencak and Ruf [82] indicated MUFA levels were highest in elephant (*Loxodonta africana*) (39%), domestic sheep (*Ovis aries*) (40%), and domestic cow (*Bos taurus*) (35%) (Figure 7J). The closest non-domesticated relatives of cow and sheep included in the analyses, eland antelope (*Tragelaphus oryx*) and mountain sheep (*Ovis ammon*) respectively, both had much lower MUFA levels (11%), suggesting evolutionary selection for large muscle size in absence of selection for high locomotor or metabolic capabilities remarkably favored high MUFA levels in the large, slow-moving species (Figures 7K and 7L). A large proportion of the MUFA chains are expected to be in PC, especially PC 34:1 [10] (Figures 7M), and elevated expression of MBOAT2, due to either enhanced satellite cell activity or transcriptionally permissive states within mature myofibers, are leading candidate mechanisms which might mediate high MUFA levels in PC of muscle. We theorize that control of *Mboat2* expression in muscle by epigenetic- and/or satellite cell-controlled mechanisms may be evolutionarily determined in various species to balance athletic performance advantages of PUFA-rich membranes to the antioxidant and rejuvenative advantages of MUFA-rich membranes. Similar pressures and mechanisms may also be active in muscle during developmental growth as well as during regeneration, and might also apply to the increased PC 34:1 and decreased PC 34:2 levels reported to occur in muscles of brown bears during hibernation [83].

The current study offers a new conceptual framework which may advance membrane-based theories of aging, and suggests that by leveraging the remarkable plasticity of MBOAT2 and MUFA-levels in PC levels of muscle, future medicines may provide age-resistant and rejuvenative qualities inherent to newborn muscle to aged muscles, possibly with tradeoffs in athletic capacities that are outweighed by therapeutic gains. Enhanced *Mboat2* expression and PC 34:1 levels in muscle might be achievable by several strategies (Figure 7N) including (1) satellite cell stimulation or (2) *Mboat2* transgene delivery. Epigenetic derepression (3) is also a promising strategy for future medicines, as *Mboat2* was among the most progressively silenced and hypermethylated loci during several stages of bovine muscle development [74]. Such medicines may ameliorate oxidative damage that drives muscular dystrophy, muscle atrophy, and sarcopenia, and might slow or reverse muscle aging.

## Acknowledgements

We thank Katsuyuki Nagata (Jikei University) for excellent technical support and critical reading of the manuscript, Fumie Hamano (University of Tokyo) for excellent support for GC-FID analyses, and Keizo Waku (Teikyo University) for invaluable advice and guidance. This study was supported by a grant from the Mishima Kaiun Foundation (WJV), the Japan Agency for Medical Research and Development (AMED) under grant numbers 22gm0910011 (HS), the NCGM Intramural Research Fund 22T001(HS), and ONO Medical Research Foundation (HS). Department of Lipid Life Science, Japan Institute for Health Security, collaborates with ONO Pharmaceutical Company Limited (Osaka, Japan) and Shimadzu Corporation (Kyoto, Japan); however, both companies did not financially support this manuscript. Department of Lipidomics, University of Tokyo, receives support from Shimadzu Corporation; however, this company had no role in the study design, data collection and analysis, decision to publish, or preparation of the manuscript. This work was also supported by a Grant-in-Aid from the Japan Society for the Promotion of Science (JSPS) KAKENHI (YA: 21H02843 and 23K21412) and an Intramural Research Grant (YA: Grant numbers 2–6 and 5-7) from National Center of Neurology and Psychiatry.

## STAR METHODS

### RESOURCE AVAILABILITY

### Lead contact

Requests for further information and resources should be directed to and will be fulfilled by the lead contact, William J. Valentine.

### Materials availability

This study did not generate new unique reagents.

### Data and code availability

GEO data sets analyzed in this study are publicly available. Accession numbers are listed in the key resources table.

All data reported in this paper will be shared by the lead contact upon request.

This paper does not report original code.

Any additional information required to reanalyze the data reported in this paper is available from the lead contact upon request.

## EXPERIMENTAL MODEL AND SUBJECT DETAILS

### Mouse experiments

All animal experiments were in accord with the Public Health Service (PHS) Policy on Humane Care and Use of Laboratory Animals and were approved and performed following the guidelines of the Animal Research Committee of the National Center for Global Health and Medicine, Japan and the Experimental Animal Care and Use Committee of the National Institute of Neuroscience of the National Center of Neurology and Psychiatry, Japan.

*Dystrophin* is an X-chromosome-linked gene and DMD affects primarily males; therefore, all mouse tissues used for analyses in this study were from male mice, except where explicitly stated. All tissues used in comparative analyses of WT background strains versus *Lpcat1*-KO, *Mboat2*-KO, and/or *mdx* were obtained from mixed cohorts of littermate mice born to parents heterozygous for the respective null alleles. Mice were housed under specific pathogen-free facilities under 12-h light/dark cycles and had free access to food and water. All mice were fed CE-2 standard rodent diet (CLEA Japan, Inc., Tokyo, Japan), except for mice raised on test diets where specified. Test diets rich in 18:1 or 18:2 also utilized and described in our previous study [34] were obtained from Research Diets, Inc. (New Brunswick, NJ) and were fed to mice from birth starting with the nursing mothers. Nutritional contents are listed in Tables S3 and S4.

Wild-type C57BL/6, B10-ScN and B10-ScSn-Dmd (mdx) mice were purchased from CLEA Japan, Inc. (Tokyo, Japan). B10-ScN, and B10-ScSn-Dmd (mdx) were backcrossed and female mice heterozygous for the *mdx* mutation were bred to produced mixed litters containing both B10-WT and B10-*mdx* male mice. A recessive mutation of *Tlr4* in the original B10-ScN strain [87] was monitored during the backcrossings, and only mice with non-mutated *Tlr4* alleles, including matched cohorts of *Tlr4*-heterozygous mice, were analyzed; other tissues from these same cohorts of mice were utilized in our previous study [34].

*Lpcat1* global gene-deficient mice (*Lpcat1*-KO) were described previously [56] and maintained on the C57BL/6 background. *Lpcat1*-KO mice were backcrossed with wild-type C57BL/6 N mice. Mice heterozygous for the *Lpcat1*-mull allele were bred and *Lpcat1*-WT and -KO male mice born to to these litters were used for comparative analyses of WT and *Lpcat1*-KO tissues.

*Mboat2* global gene-deficient (*Mboat2*-KO) mice were generated on the C57BL/6 J background by CRISPR-Cas9–mediated genome editing by Department of Laboratory Animal Medicine at Japan Institute for Health Security as described previously [88]. Briefly, the crRNA guiding sequence for the exon 10 of *Mboat2* gene was as follows: 5ʹ-TGTCTGGATTGTCGGACTGA -3ʹ. Both crRNA and tracrRNA were chemically synthesized (Fasmac, Kanagawa, Japan). The recombinant Cas9 protein (30ng/μl, New England Biolabs, MA, USA), crRNA (12.2 pmol/μl) and tracrRNA (12.2 pmol/μl) were co-injected into the cytoplasm of pronuclear stage eggs from C57BL/6JJcl mice (CLEA Japan, Inc., Tokyo, Japan). After overnight culture, two-cell stage embryos were transferred into the oviduct of pseudopregnant ICR mice anesthetized using the mixture of 0.3 mg/kg medetomidine, 4.0 mg/kg midazolam, and 5.0 mg/kg butorphanol during operation [89]. The mutation in *Mboat2* was confirmed by Sanger sequencing analysis. *Mboat2*-KO mice on C57BL/6 background were transferred to the National Center of Neurology and Psychiatry and backcrossed with *mdx* mice on the C57BL/6J background (B6-*mdx*) to produce *Mboat2*-gene deficient *mdx* (*Mboat2*-KO/*mdx*) mice. For dystrophic *mdx* and muscle injury models, mice heterozygous for the *Mboat2*-null and *mdx* mutant alleles were bred to produce mixed litters containing all combinations of WT, *mdx*, *Mboat2*-KO, and *Mboat2*-KO/*mdx* mice.

### Genotyping

Mouse genotyping of *mdx* and *Mboat2* status was performed from by primer competition PCR [90]. The three-primer set to detect *mdx* status was common-forward (5’-GCG CGA AAC TCA TCA AAT ATG CGT GTT AG TGT-3’), WT-reverse (5’-GAT ACG CTG CTT TAA TGC CTT TAG TCA CTC AGA TAG TTG AAG CCA TTT TG-3’), and *mdx*-reverse (5’-CGG CCT GTC ACT CAG ATA GTT GAA GCC ATT TTA-3’), which detected the WT (134 base pairs) and mutant alleles (117 base pairs). The three-primer set to detect *Mboat2* status was common-forward (5’-GCG CGT TCT CTT GGC TCT CCT GTT ATC TAT G-3’), WT-reverse (5’-GAT ACG CTG CTT TAA TGC CTT TAA GAA AGA ATG TCT GGA TTG TCG GAC T-3’), and mutant-reverse (5’-CGG CCA GAA AGA ATG TCT GGA TTG TCG GAA G-3’), which detected WT (113 base pairs) and mutant alleles (94 base pairs). *Lpcat1* genotyping was performed by standard PCR amplification using forward (5’-ATT AAC CAC TGA CAA GCA GGT GCT C-3’) and reverse (5’-TCA TGC TAC AGG GAA CCA CAG TGA C-5’) primers, which detected WT (475 base pairs) and mutant (184 base pairs) alleles.

### Research test diets

Mouse test diets rich in 18:1 or 18:2, also utilized and described in our previous study [34], were obtained from Research Diets, Inc. (New Brunswick, NJ) and were fed to mice from birth starting with the nursing mothers. Nutritional contents are listed in Tables S3 and S4.

### Muscle injury

Mice were anesthetized by isoflurane inhalation before intramuscular injections barium chloride or cardiotoxin solutions, as indicated in figure legends, to induce muscle injury [59]. Right TA muscles were injected with 30 µL of 1.2% (w/v) barium chloride solution in saline for barium chloride injury and 30 µL of a 10 µm cardiotoxin solution for cardiotoxin injury. Left, uninjected TA muscles were controls. At indicated time points, mice were euthanized by isoflurane inhalation followed by cervical dislocation, and muscles were harvested for analysis.

## METHOD DETAIL

### C2C12 cell culture

C2C12 cells were cultured at 37°C in a humidified atmosphere of 5% CO_2_ in growth media consisting of DMEM (Gibco) supplemented with 10% heat-inactivated (56°C, 30 min) FBS (Gibco) and 2 mM glutamine. Differentiation was induced by culturing in differentiation media consisting of DMEM containing 2% heat-inactivated horse serum (Gibco) and 2 mM glutamine, the media was replenished for 7 days, then myotubes were selectively enriched from undifferentiated cells based on adherence properties [91, 92]. For myotube enrichment, the myotube cell cultures were detached in trypsin/EDTA (0.05%/0.002%) and then pre-plated on 10 cm culture dishes for 30–60 min in growth media to remove the more rapidly adhering undifferentiated cells. Culture media containing unattached myotubes was collected and centrifuged (240 x *g*, 5 min). The pelleted myotubes were resuspended in differentiation media and plated onto 24-well Corning tissue culture pre-coated with 0.1% gelatin, and undifferentiated C2C12 cells were plated in growth media (20,000 cells/well). Cell extracts were collected two days later for analysis by RT-PCR.

### CHO-K1 cell culture

CHO-K1 cells were cultured at 37°C in a humidified atmosphere of 5% CO_2_ in growth media consisting of Ham’s F12 media (Nacalai Tesque) supplemented with 10% FBS.

### Isolation and culture of mouse primary satellite cells by pre-plate method

Pre plate method was used for isolation of all primary satellite cells, except for comparative analyses of *Mboat2*-WT and -KO satellite cells, noted below. For primary WT satellite cell cultures, cells were isolated from hind limbs of adult (4-week-old to one-year-old) male and female C57BL/6 mice using modified method of Rando and Blau [51, 93]. Hind limb muscles were minced in PBS supplemented with 5 mM CaCl_2_, 5 mM collagenase, and dispase I (400 protease units / ml) and incubated at 37°C for 1-2 h and intermittently triturated. The digested slurry was diluted with PBS and passed through 100 μm and then 40 μm cell strainers (Falcon). The filtrate was centrifuged at 350 x *g* for 15 min and the pelleted cells were resuspended in growth media consisting of Ham’s F-10 media (Gibco) supplemented with 20% heat-inactivated FBS, 1% anti-biotic anti-mycotic (Sigma), and 5 ng/ml basic fibroblast growth factor. The cells were cultured on petri dishes (Rikaken, RSU-SD9015-2) coated with type I collagen (Sigma, C8919). Culture medias were replaced every two or three days, and cells were maintained at less than 70% confluency. Satellite cells were enriched during early passages by detachment with PBS followed by pre-plating [94] and used for experiments within ten passages. To induce differentiation, satellite cells were seeded onto type I collagen-coated 24-well plates (Iwaki, 4820-010) in growth media and the next day growth media was replaced with differentiation media consisting of DMEM containing 2% heat-inactivated horse serum, 1% chick embryonic extract ultrafiltrate (US Biological Life Sciences), and 2 mM glutamine. Differentiation media was replenished daily. Where indicated in the text, medias were supplemented with a fatty acid mixture (5 μM each of FA 18:2, FA 20:4, and FA 22:6) to supply additional substrates for phospholipid production.

### FACS cell-sorting of *Mboat2*-WT and *Mboat2*-KO satellite cells

FACS sorting was performed only for comparative analyses of *Mboat2*-WT and -KO satellite cells, using modified methods of previous reports [95, 96]. Hind limb mouse muscles were isolated and minced in a DMEM containg 10% FBS, 3.5 mg/mL Type 2 Collagenase (Worthington), and dispase I (400 protease units / ml) and incubated at 37°C for 1-2 h and intermittently triturated. The digested slurry was diluted with PBS and passed through 100 μm and then 40 μm cell strainers (Falcon). The filtrate was centrifuged at 350 x *g* for 15 min and the pelleted cells were resuspended in DMEM supplemented with 10% FBS and stained with anti-CD31-FITC antibody (1:200; clone 390; eBioscience), anti-CD45-FITC antibody (1:200; clone 30-F11; eBioscience), anti-Sca1-PE antibody (1:200; clone D7; eBioscience), anti-integrin α7 antibody (1:200; clone 3C12; MBL International, Woburn, MA, USA), and anti-mouse IgG Alexa Fluor 647 antibody (1:200; Jackson Immuno Research Laboratory)[95]. After staining, satellite cells were isolated by FACS using a SONY SH800S flow cytomoter. Satellite cells were cultured in growth medium consisting of DMEM supplemented with 20% FBS, 1% anti-biotic anti-mycotic (ABAM), and 5 ng/ml basic fibroblast growth factor on culture dishes coated with growth-factor-reduced Matrigel. Medium was replaced with DMEM containing 2% horse serum to induce differentiation.

### LPCAT substrate choice assay

LPCAT substrate choice biochemical assays [53] were performed similar to our previous studies [54, 56]. CHO-K1 cells were transfected with expression plasmids for *Mboat2* or vector (pCDNA3.1) using Lipofectamine L2000 acccording to the manufacturer’s instructions and cells were harvested after24 h. Microsomal membrane fractions were prepared by sonication of cells on ice using a probe sonicator (Ohtake Works) for 3 x 30 seconds in TSC buffer: 100 mM Tris-HCl (pH 7.4), 300 mM sucrose, 1 mM EDTA, and 1X complete protease inhibitor cocktail (Roche). Sonicated samples were centrifuged at 9000 x *g* for 10 minutes to remove nuclei and debris, then supernatants were ultracentrifuged at 100,000 x *g* for 1 hour. Pellets were resuspended in TSC and snap frozen in aliquots. Protein concentrations were measured using a Pierce BCA Protein Assay Kit. LPCAT activity was measured by incubating membrane fractions (containing 0.01 μg of protein) with substrates at 37°C for 10 minutes in 100 µL reaction mixtures containing 25 μM deuterium-labeled LPC 16:0 and 1 μM each 16:0-, 18:1-, 18:2-, 20:4-, and 22:6-CoA (Avanti). The reaction buffer contained 110 mM Tris-HCl (pH 7.5), 1 mM EDTA, 2 mM CaCl_2_, and 0.015% Tween-20. Reactions were halted by addition of 375 µL methanol/chloroform (2:1). Dilauryl-PC (Avanti) was added as an internal standard, lipids were extracted by Bligh/Dyer method [97], dried using a centrifugal evaporator (Sakuma Seisakusho Ltd., Tokyo, Japan), and resuspended in methanol. Products were measured by LC-MS using a TSQ Vantage triple stage quadrupole mass spectrometer (Thermo Fisher Scientific) by selected reaction monitoring (SRM) using the transition [M + H]+ → 184.1 to detect PC (positive ion mode electrospray ionization). Signals of LPCAT products were compared to calibration curves of nonlabelled standards for quantification and normalized to the internal standard.

### siRNA transfection of satellite cells

ON-TARGET plus siRNAs targeting murine *Mboat2*, *Agpat3*, and control siRNA were purchased from Dharmacon. The siRNAs used were J-063482-10 (si*Mboat2*#2), J-063482-12 (si*Mboat2*#1), J-059804-12 (si*Agpat3*), and D-001810-10-20 ON-TARGETplus non-targeting Pool (siControl). Satellite cells were plated onto 24-well collagen-coated tissue culture plates (Iwaki) in growth media (150,000 cells/well) the day before siRNA transfection. The next day, culture medias were changed to differentiation media and siRNA transfection complexes were applied to each well. Transfection complexes consisting of 1.5 µL RNAiMAX and 5 pmole siRNA complexed in 100 µL of DMEM.

### RT-PCR

Total RNA was isolated from C2C12 cells and satellite cells using RNeasy Mini Kits and from tissues also using RNeasy fibrous tissue Kits (Qiagen, Valencia, CA, United States). cDNAs were synthesized using random primers and SuperScript III reverse transcriptase (Invitrogen) or High-Capacity cDNA Reverse Transcription kit (ThermoScientific). RT-PCR analysis was performed using a StepOnePlus Real-Time PCR System and Fast SYBR Green Master Mix (Applied Biosystems). mRNA expression levels were normalized to *18S* rRNA and fold-changes were calculated using the 2^−ΔΔCT^ method. The PCR primer sequences are listed in Table S6.

### Lipid extraction from cells and tissues

Lipid extractions from cells and tissues were performed as recentlydescribed [98] (in press). In brief, satellite cells were cultured on 24 well plates, and cellular phospholipids were extracted using methanol (500 µL/well) [99]. After incubation for 5 min at room temperature, the extracts were centrifuged at 4°C for 10 min at 18,700 x *g*, and supernatants were collected and further diluted in methanol before LC-MS analysis.

Whole muscles or tissue pieces from mid-belly regions of muscles were flash frozen in liquid nitrogen and stored at -80°C. Tissue pieces were pulverized in a frozen state using a Tokken Automill cryogenic pulverizer (Tokken, Japan). Whole muscles were cryosectioned and frozen sections (10–20 µm in thickness) were obtained from the mid-belly regions. Methanol was added to the frozen crushed tissue pieces or muscle sections (3–25 µg) and the samples were rocked end-over-end for 1 h at 4°C and then centrifuged at 18,700 x *g* for 10 min at 4 C. The supernatants were collected, diluted with methanol to a concentration corresponding to 3 mg of original tissue/ml, and stored at 80°C.

### Lipidomic analyses

Samples were further diluted with methanol ten-fold before LC-selected reaction monitoring (SRM)-MS analyses were performed using a Nexera UHPLC system and LC-MS using Shimadzu 8050 triple quadrupole mass spectrometers (Shimadzu, Japan). An Acquity UPLC BEH C8 column (1.7 μm, 2.1 mm x 100 mm, Waters) was used with the following ternary mobile phase compositions: 5 mM NH_4_HCO_3_/water (mobile phase A), acetonitrile (mobile phase B), and isopropanol (mobile phase C). The pump gradient [time (%A/%B/%C)] was programmed as follows: 0 min (50/45/5)-10 min (20/75/5)-20 min (20/50/30)-27.5 min (5/5/90)-28.5 min (5/5/90)-28.6 min (50/45/5). The flow rate was 0.35 ml/min and the column temperature was 47°C. The injection volume was 5 μl.

Comprehensive LC-SRM-MS analysis was performed in the positive ion mode electrospray ionization with the transitions [M+H]+ →184 for PC and [M+H]+ →[M+H-141]+ for PE to detect all diacyl PC and PE species possessing even number carbon chains 14–24 carbons in length in sn-1 and sn-2. Chromatographic peak areas were used for comparative quantitation of each molecular species (e.g., 38:6) within a given class (e.g., PC). Peak areas were calculated by Traces software [100]. Peak areas of each individual species were normalized against the sum of all peak areas within that class to determine the relative abundances (expressed as % of total), and all major species comprising at least 3% of total PC or PE signals were plotted, except in figures 1D and 1F where a 5% threshold was set. Plasmalogen may be abundant components of PE and generally exist at low levels in PC; however, the methodologies we employed are inefficient to detect plasmalogen species [101]. Therefore, we limited our analyses to diacyl phospholipids, and all signals assigned to plasmalogen species were omitted from our calculations.

### Muscle fiber diameter measurements

Transverse sections (10 µm) were cut from the mid-belly regions of TA muscles which had been flash frozen in liquid nitrogen-chilled isopentane. Sections were stained by hematoxylin and eosin. TA muscle sections were imaged and analyzed using a Keyence BZ-X800 Image system. The minimum diameters of at least 200 muscle fibers per muscle were measured. The variance coefficient of the muscle fiber diameters were determined by the equation: variance coefficient = (standard deviation of the muscle fiber diameter / mean muscle fiber diameter) X 1000 [102].

### AAV-*Mboat2* expression

AAV-expression particles under control of human skeletal muscle actin alpha 1 (ACTA1) were purchased from Vector Builder to express coding regions for eGFP (AAV-*eGFP*), mouse *Mboat2* (AAV-*Mboat2*), or an enzymatically inactive mutant of *Mboat2* (AAV-inactive mutant) with a (WH>AA) mutation in the conserved MBOAT Motif B which is essential for LPLAT activity [11]. Enzymatic activity of the AAV-*Mboat2* transgene and full lack of activity for the AAV-inactive mutant *Mboat2* trangene were confirmed in LPCAT substrate choice biochemical assays (data not shown). The transgene were packaged into recombinant AAV serotype 1 capsids, which are efficient to transduce mouse muscle by intramuscular injection [103]. In initial experiments, escalating doses (2 x 10^9^ - 2 x 10^11^) of AAV-*Mboat2* particles were injected in one gastrocnemius muscle and of AAV-*eGFP* were injected in the other.

## QUANTIFICATION AND STATISTICAL ANALYSES

All data plotted in bar graphs represent means. Statistical significance were determined by t-tests (two-tailed) or Bonferroni’s multiple-comparison tests, as indicated in figure legends. *P* values < 0.05 were considered statistically significant. (*p< 0.05, **p< 0.01, ***p< 0.001; ns, not significant). Sample size for each experiment is indicated in the figure legend. RT-PCR expression values were log transformed before statistical analyses [104]. All statistical analyses were performed using GraphPad Prism10 (GraphPad Software, La Jolla, CA).

## STAR METHODS

### Key resources table

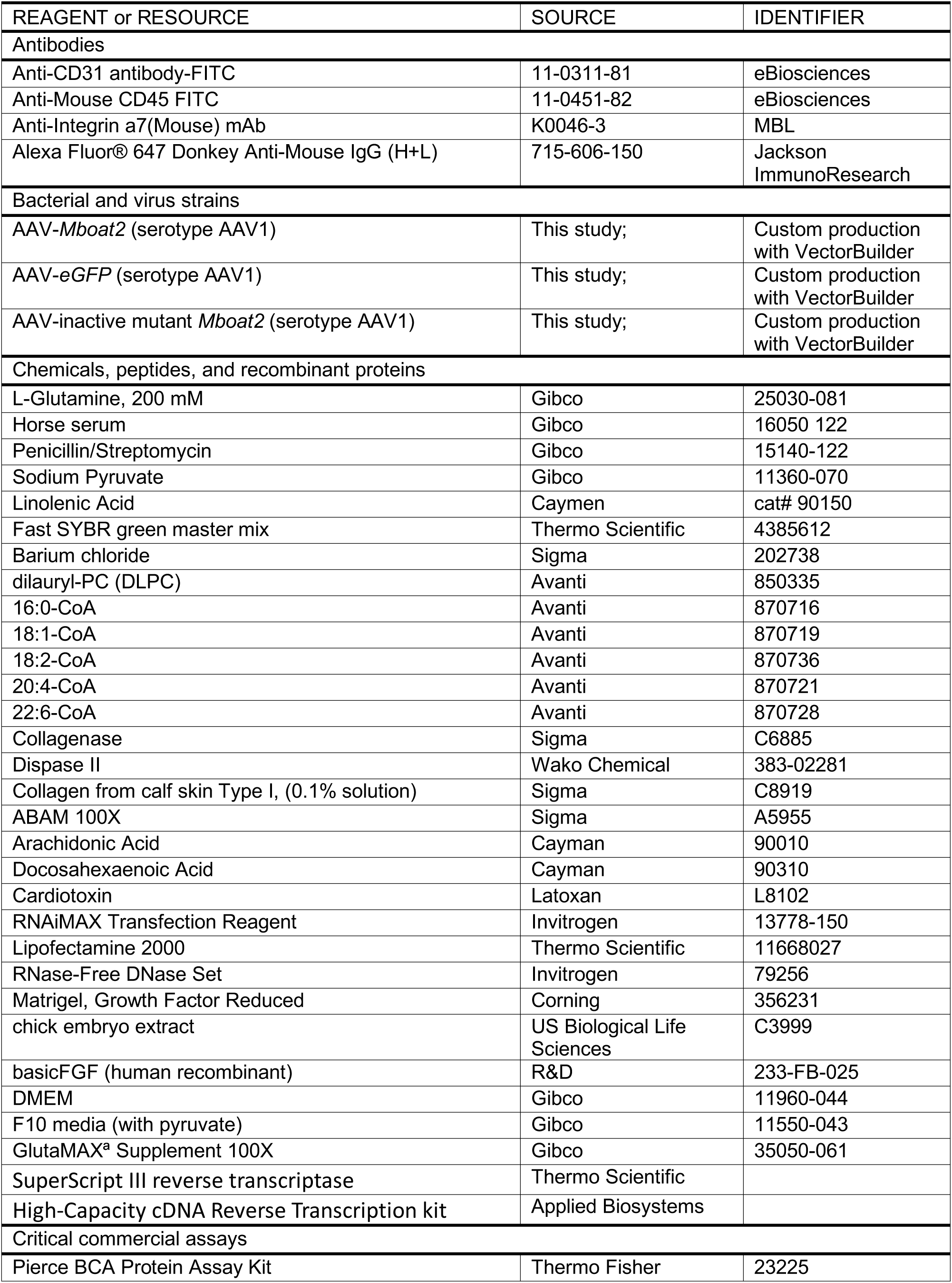
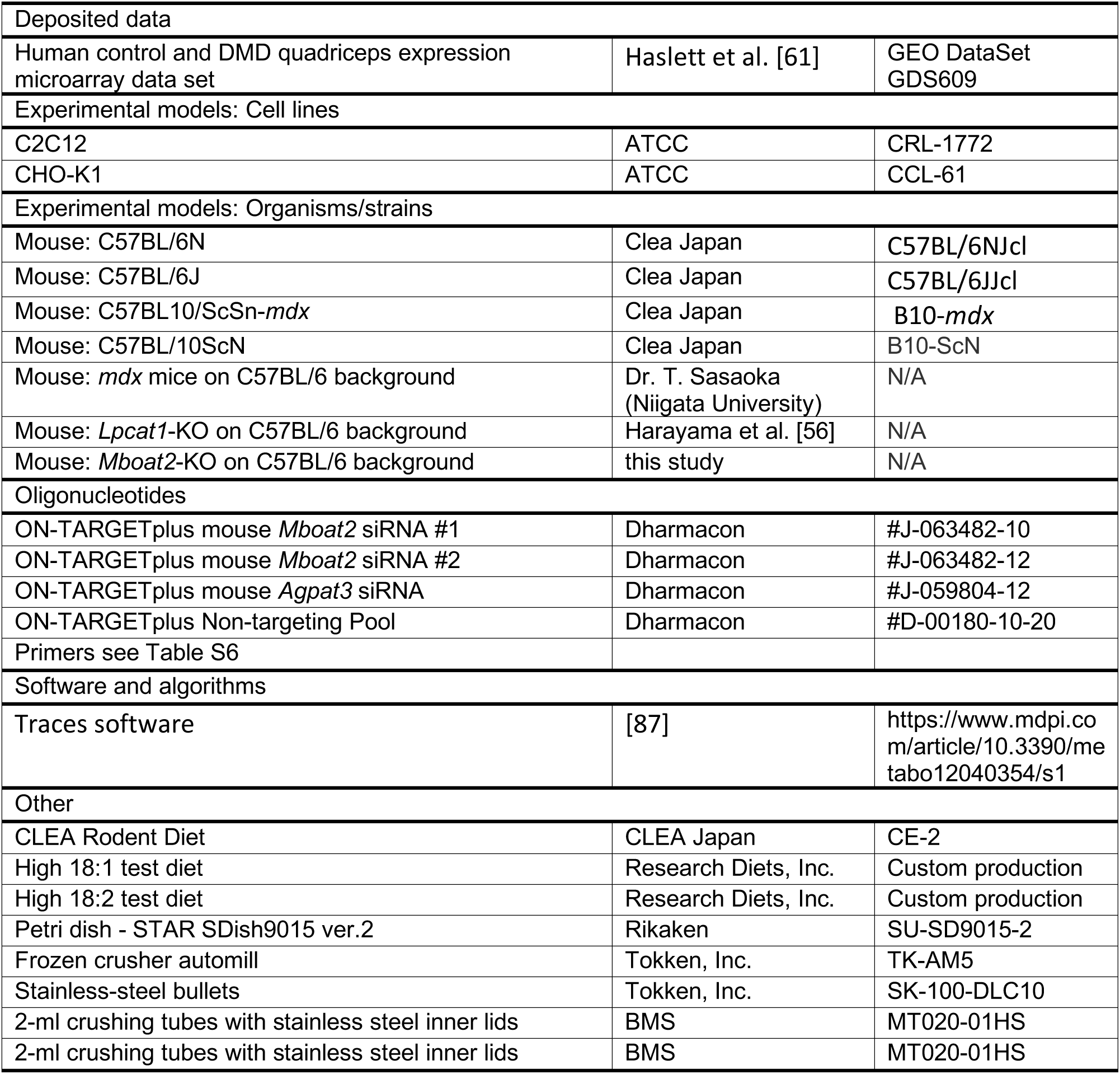

