## supplemental for "MBOAT2 limits the amounts of PUFA in phosphatidylcholine of neonatal and dystrophic skeletal muscle and promotes muscle regeneration"

### Figure S1

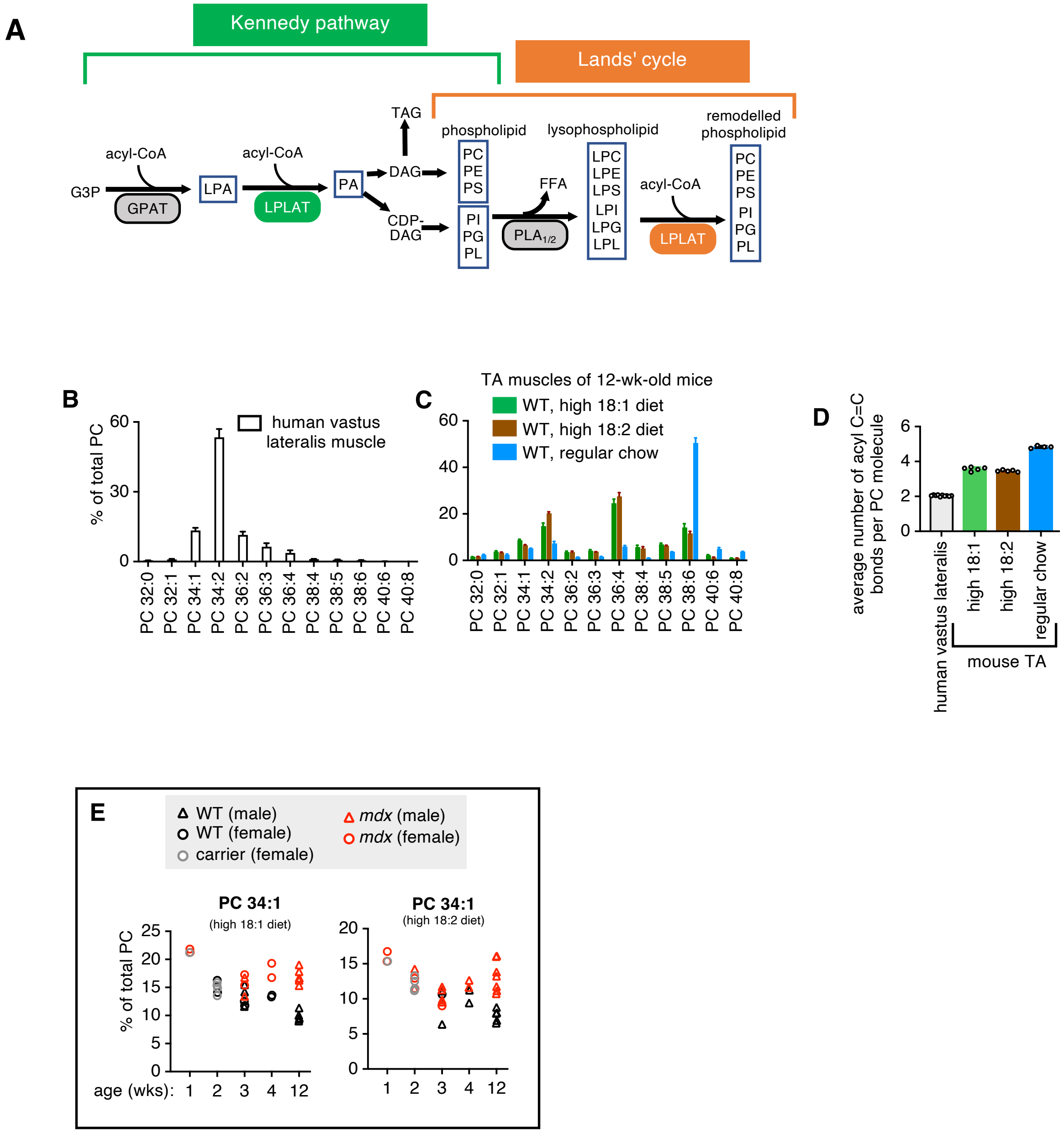
A
Kennedy pathway
Lands' cycle
G3P
acyl-CoA
GPAT
LPA
acyl-CoA
LPLAT
PA
TAG
DAG
CDP-DAG
phospholipid
PC PE PS
PI PG PL
FFA
PLA1/2
lysophospholipid
LPC LPE LPS LPI LPG LPL
acyl-CoA
LPLAT
remodelled phospholipid
PC PE PS PI PG PL
B
% of total PC
human vastus lateralis muscle
C
TA muscles of 12-wk-old mice
WT, high 18:1 diet
WT, high 18:2 diet
WT, regular chow
D
average number of acyl C=C bonds per PC molecule
human vastus lateralis
high 18:1
high 18:2
regular chow
mouse TA
E
WT (male)
WT (female)
carrier (female)
mdx (male)
mdx (female)
PC 34:1
(high 18:1 diet)
PC 34:1
(high 18:2 diet)
% of total PC
age (wks):

### Figure S2

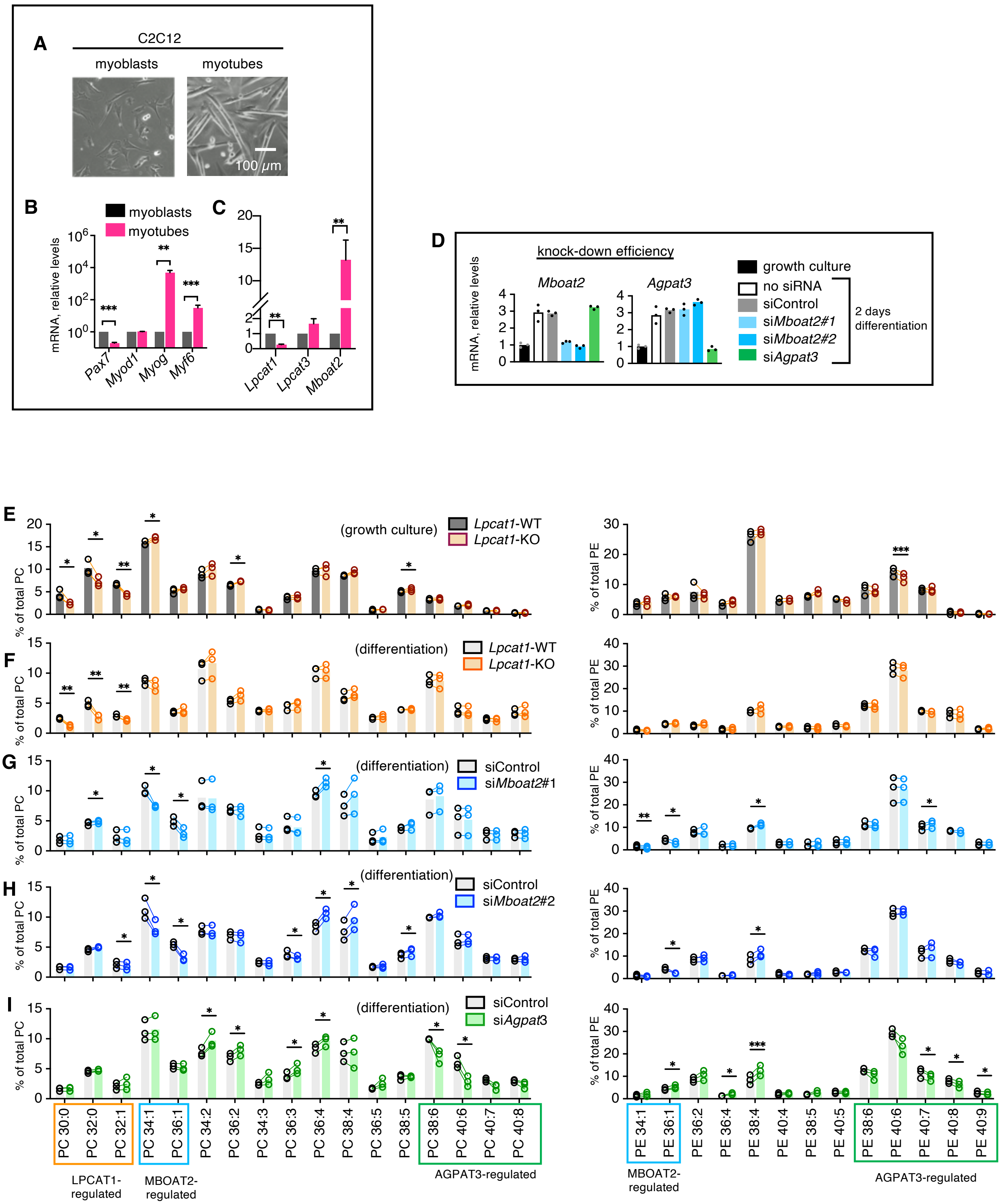
A
C2C12
myoblasts
myotubes
100 μm
B
myoblasts
myotubes
mRNA, relative levels
Pax7
Myod1
Myog
Myf6
C
Lpcat1
Lpcat3
Mboat2
D
knock-down efficiency
Mboat2
Agpat3
growth culture
no siRNA
siControl
siMboat2#1
siMboat2#2
siAgpat3
2 days differentiation
E
(growth culture)
Lpcat1-WT
Lpcat1-KO
% of total PC
% of total PE
F
(differentiation)
G
siControl
siMboat2#1
H
siMboat2#2
I
siAgpat3
PC 30:0
PC 32:0
PC 32:1
PC 34:1
PC 36:1
PC 34:2
PC 36:2
PC 34:3
PC 36:3
PC 36:4
PC 38:4
PC 36:5
PC 38:5
PC 38:6
PC 40:6
PC 40:7
PC 40:8
LPCAT1-regulated
MBOAT2-regulated
AGPAT3-regulated
PE 34:1
PE 36:1
PE 36:2
PE 36:4
PE 38:4
PE 40:4
PE 38:5
PE 40:5
PE 38:6
PE 40:6
PE 40:7
PE 40:8
PE 40:9

### Figure S3

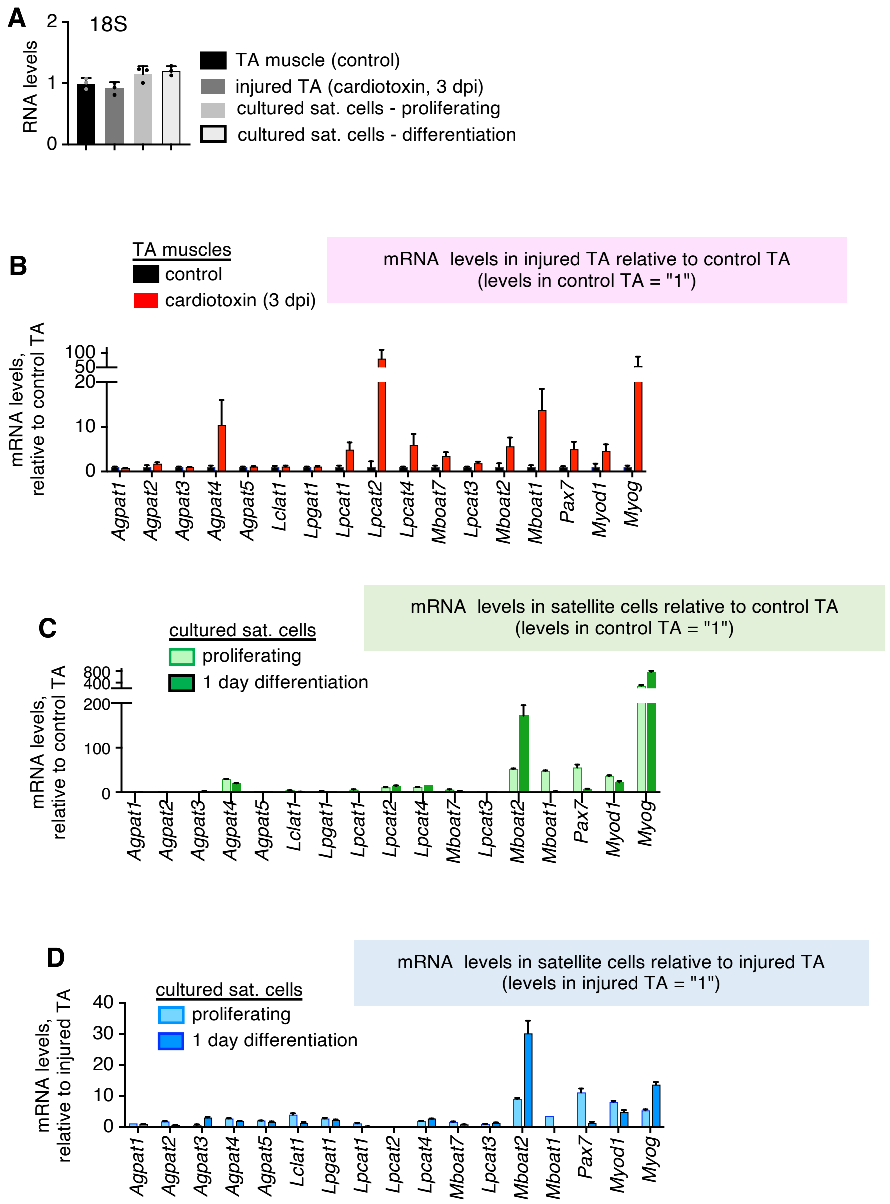
A
18S
RNA levels
2
1
0
TA muscle (control)
injured TA (cardiotoxin, 3 dpi)
cultured sat. cells - proliferating
cultured sat. cells - differentiation
B
TA muscles
control
cardiotoxin (3 dpi)
mRNA levels in injured TA relative to control TA
(levels in control TA = "1")
mRNA levels,
relative to control TA
100
50
20
10
0
Agpat1
Agpat2
Agpat3
Agpat4
Agpat5
Lclat1
Lpgat1
Lpcat1
Lpcat2
Lpcat4
Mboat7
Lpcat3
Mboat2
Mboat1
Pax7
Myod1
Myog
C
cultured sat. cells
proliferating
1 day differentiation
mRNA levels in satellite cells relative to control TA
(levels in control TA = "1")
mRNA levels,
relative to control TA
800
400
200
100
0
Agpat1
Agpat2
Agpat3
Agpat4
Agpat5
Lclat1
Lpgat1
Lpcat1
Lpcat2
Lpcat4
Mboat7
Lpcat3
Mboat2
Mboat1
Pax7
Myod1
Myog
D
cultured sat. cells
proliferating
1 day differentiation
mRNA levels in satellite cells relative to injured TA
(levels in injured TA = "1")
mRNA levels,
relative to injured TA
40
30
20
10
0
Agpat1
Agpat2
Agpat3
Agpat4
Agpat5
Lclat1
Lpgat1
Lpcat1
Lpcat2
Lpcat4
Mboat7
Lpcat3
Mboat2
Mboat1
Pax7
Myod1
Myog

### Figure S4

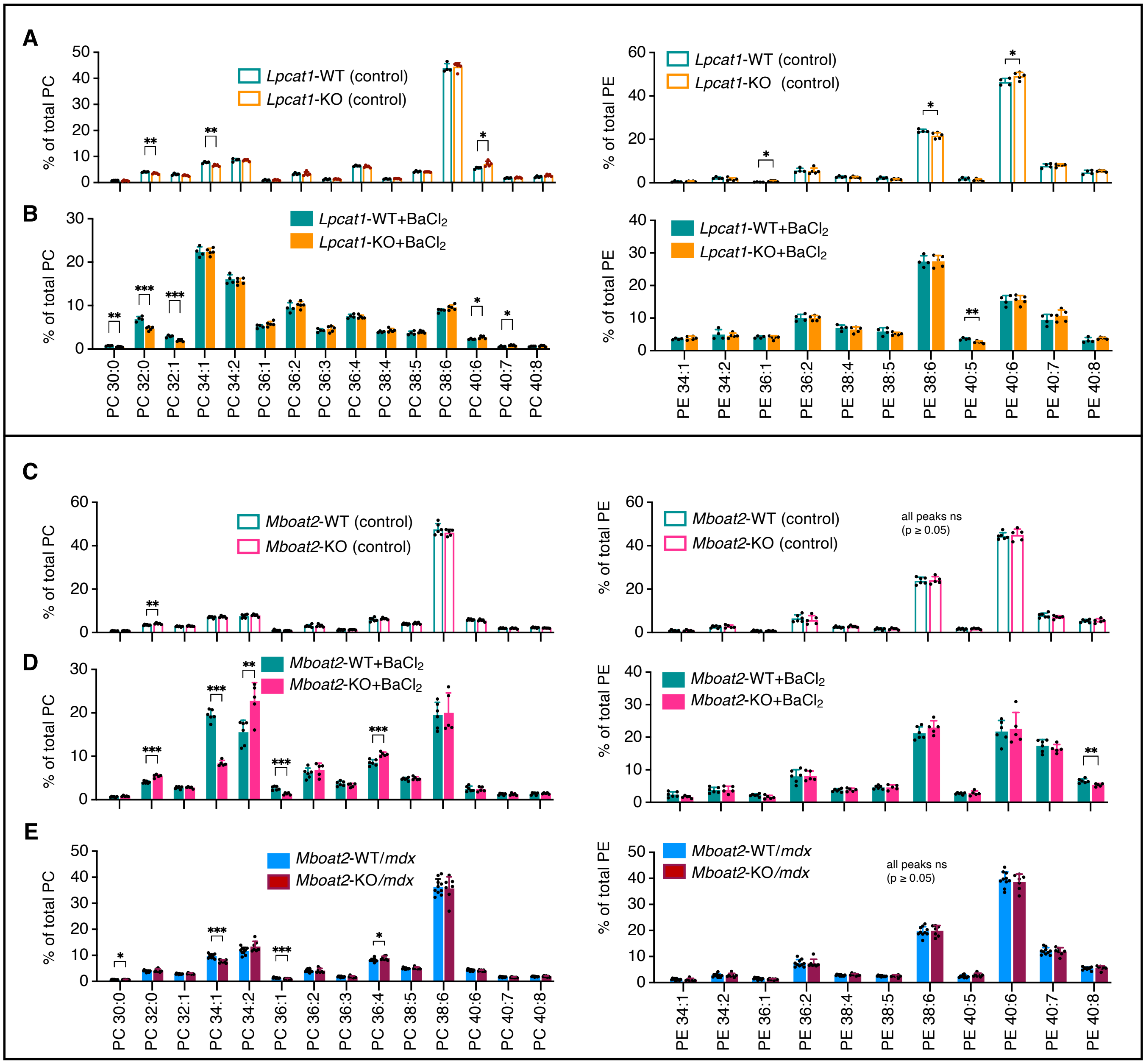

A
Lpcat1-WT (control)
Lpcat1-KO (control)
% of total PC
% of total PE
B
Lpcat1-WT+BaCl2
Lpcat1-KO+BaCl2
C
Mboat2-WT (control)
Mboat2-KO (control)
all peaks ns
(p ≥ 0.05)
D
Mboat2-WT+BaCl2
Mboat2-KO+BaCl2
E
Mboat2-WT/mdx
Mboat2-KO/mdx
PC 30:0 PC 32:0 PC 32:1 PC 34:1 PC 34:2 PC 36:1 PC 36:2 PC 36:3 PC 36:4 PC 38:4 PC 38:5 PC 38:6 PC 40:6 PC 40:7 PC 40:8
PE 34:1 PE 34:2 PE 36:1 PE 36:2 PE 38:4 PE 38:5 PE 38:6 PE 40:5 PE 40:6 PE 40:7 PE 40:8

### Figure S5

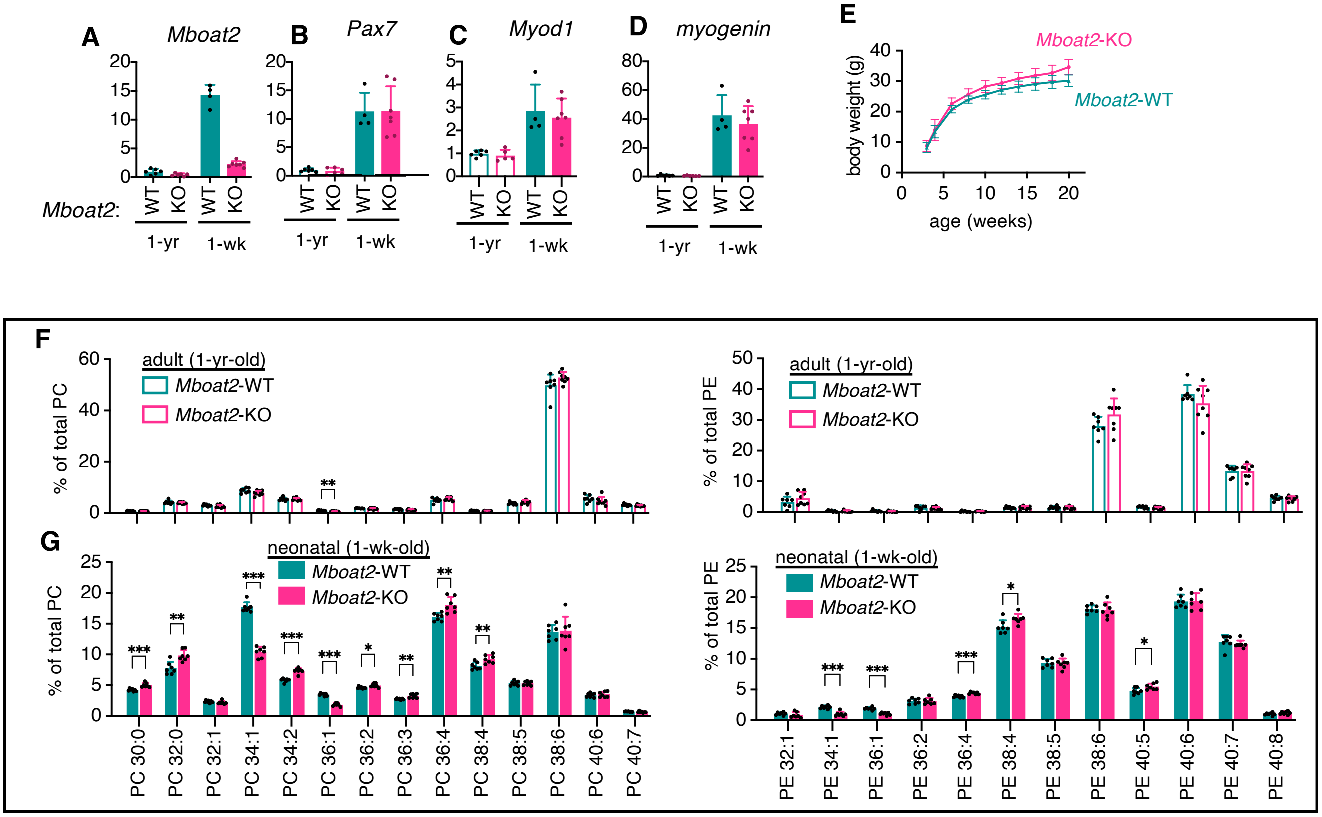

A
Mboat2
B
Pax7
C
Myod1
D
myogenin
Mboat2:
WT
KO
1-yr
1-wk
E
Mboat2-KO
Mboat2-WT
body weight (g)
age (weeks)
F
adult (1-yr-old)
Mboat2-WT
Mboat2-KO
% of total PC
% of total PE
G
neonatal (1-wk-old)
PC 30:0
PC 32:0
PC 32:1
PC 34:1
PC 34:2
PC 36:1
PC 36:2
PC 36:3
PC 36:4
PC 38:4
PC 38:5
PC 38:6
PC 40:6
PC 40:7
PE 32:1
PE 34:1
PE 36:1
PE 36:2
PE 36:4
PE 38:4
PE 38:5
PE 38:6
PE 40:5
PE 40:6
PE 40:7
PE 40:8

### Figure S6

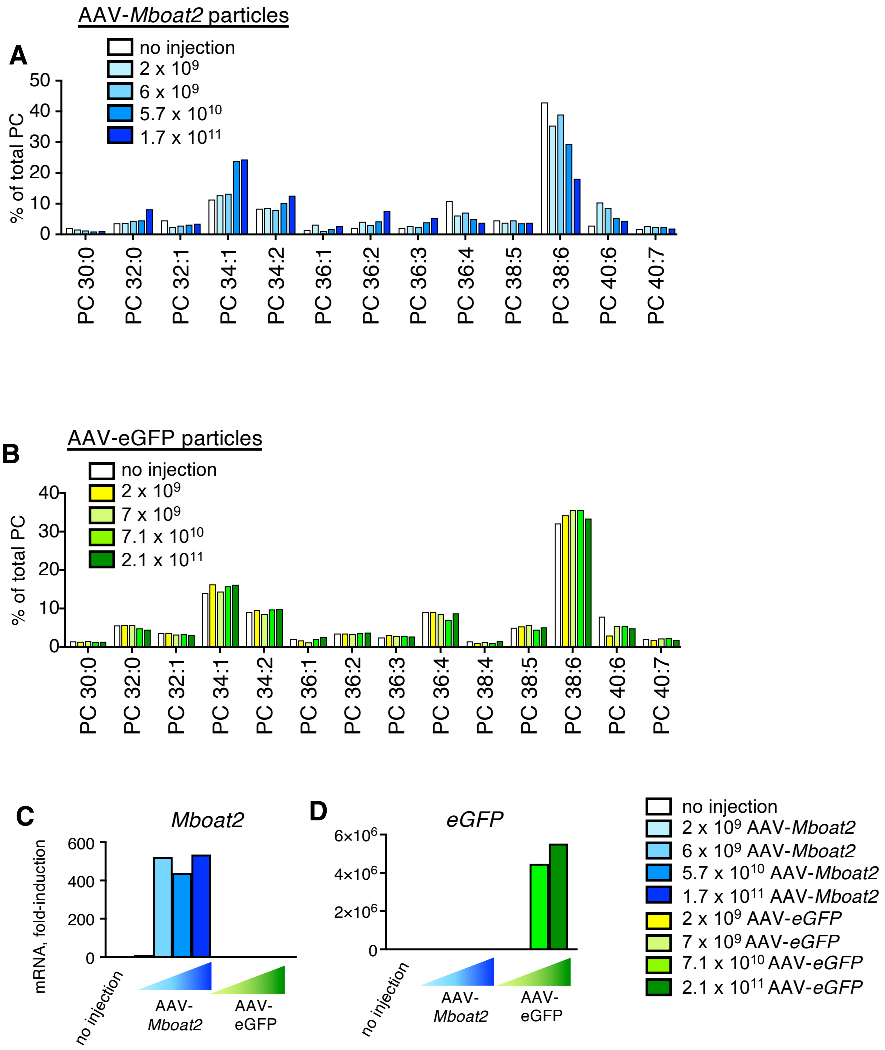
A
AAV-Mboat2 particles
no injection
2 x 10^9
6 x 10^9
5.7 x 10^10
1.7 x 10^11
% of total PC
PC 30:0
PC 32:0
PC 32:1
PC 34:1
PC 34:2
PC 36:1
PC 36:2
PC 36:3
PC 36:4
PC 38:5
PC 38:6
PC 40:6
PC 40:7
B
AAV-eGFP particles
no injection
2 x 10^9
7 x 10^9
7.1 x 10^10
2.1 x 10^11
% of total PC
PC 38:4
C
Mboat2
mRNA, fold-induction
no injection
AAV-Mboat2
AAV-eGFP
D
eGFP
no injection
AAV-Mboat2
AAV-eGFP
no injection
2 x 10^9 AAV-Mboat2
6 x 10^9 AAV-Mboat2
5.7 x 10^10 AAV-Mboat2
1.7 x 10^11 AAV-Mboat2
2 x 10^9 AAV-eGFP
7 x 10^9 AAV-eGFP
7.1 x 10^10 AAV-eGFP
2.1 x 10^11 AAV-eGFP

#### SUPPLEMENTAL FIGURES

**Figure S1. Comparison of PC compositions of human and mouse muscle and PE compositions of dystrophic *mdx* muscle, related to Figure 1.**

(A) LPLATs regulate fatty acid compositions of phospholipids both during Kennedy pathway (*de novo* biosynthesis) and Lands' cycle remodeling. CDP-DAG, cytidine diphosphate-DAG; CL, cardiolipin; DAG, diacylglycerol; G3P, glycerol-3-phosphate; GPAT, G3P acyltransferase; LPL, lysophospholipid; LPLAT, LPL acyltransferase; PA, phosphatidic acid; PC, phosphatidylcholine; PE, phosphatidylethanolamine; PG, phosphatidylglycerol; PI, phosphatidylinositol; PLA, phospholipase; PS, phosphatidylserine; TAG, triacylglycerol.

(B) PC compositions of vastus lateralis muscle from healthy human volunteers (5 male and 5 female volunteers, age 48 to 77 years). Values for each volunteer were calculated from the average values of serial biopsies taken every 4 h across 24 h and represent data originally reported by Loizides-Mangold et al. [1].

(C) PC compositions of TA muscles of 12-week-old B10-WT mice raised on three different diets (n = 4-5 / group).

(D) Average number of acyl chain C=C bonds per PC molecule in human vastus lateralis or mouse TA muscles. Values were calculated from data plotted in panels A and B.

(E) PC 34:1 levels in gastrocnemius muscles of B10-WT and -*mdx* mice, ages 1-, 2-, 3, 4- and 12-weeks, raised on two custom diets. Individual data points are plotted. Identical to data plotted in Figure 1E, but containing additional genotype and gender information of each sample.

Error bars represent SD (B and C).

**Figure S2. LPCAT1 generates PC 32:0 and MBOAT2 generates PC 34:1 in satellite cells, related to Figure 2.**

(A-C) Representative images (A) of C2C12 cells either undifferentiated (myoblasts) or differentiated to myotubes. Expression levels of selected myogenic marker genes (B) and LPLATs (C) are plotted (n = 3 independent experiments).

(D) Related to Figures 2I-2K. Knock-down efficiencies of *Mboat2*- and *Agpat3*-targeting siRNAs. Control or gene-specific siRNAs were applied at the start of 2 days differentiation. (n = 1 experiment).

(E-I) Complete data sets that are summarized in heatmaps in Figures 2J and 2K. Satellite cells cultured were for two days in either growth media (E) or differentiation media (F-I); medias were supplemented with fatty acids (18:2, 20:4, and 22:6; 5 µM/each). *Lpcat1* gene suppression (E-F)- plotted data show comparisons between WT and *Lpcat1*-KO satellite cells in growth culture (E) or differentiation (F) conditions. *Mboat2* and *Agpat3* gene suppression (G-I)- plotted data show comparisons between WT satellite cells following two days differentiation in presence of two independent siRNAs siRNAs that target *Mboat2* (si*Mboat2*-#1 and -#2) (G and H) or a siRNA that targets *Agpat3* (si*Agpat3*) (I). Three independent experiments were performed for each gene suppression strategy.

Error bars represent SEM (B,C, E-I). Significance is based on t-tests (B,C, E-I).

**Figure S3. Expression levels of LPLATs and myogenic marker genes in primary satellite cells compared to muscle tissues, related to figure 3.**

Extended plots of data plotted in Figures 3H-3J. Comparative gene expressions in control and cardiotoxin-uninjured (3 dpi) TA muscles and primary satellite cells. LPLAT and myogenic marker gene expressions were normalized to *18S* as a reference gene in order to compare expression levels between muscle cells and tissues (n = 3 muscles/group).

(A) *18S* expression levels (not normalized) in primary satellite cells and muscle tissues using same amounts of RNA as template (levels in control TA muscles were arbitrarily set to 1).

(B) Expression levels of LPLATs and myogenic marker genes in cardiotoxin-injured TA muscles relative to the levels in control TA muscles (levels in control TA muscles were arbitrarily set to 1 following normalization to *18S*).

(C) Expression levels of LPLATs and myogenic marker genes in cultured satellite cells relative to the levels in control TA muscles (levels in control TA muscles were arbitrarily set to 1 following normalization to *18S*).

(D) Expression levels of LPLATs and myogenic markers in cultured satellite cells, relative to the levels in injured TA muscles (levels in injured TA muscles were arbitrarily set to 1 following normalization to *18S*).

**Figure S4. Comparative analyses of PC and PE compositions of the WT and KO muscle tissues, related to figure 4.**

(A-B) PC and PE compositions of control (A) or injured (barium chloride injection, 7 dpi) (B) TA muscles of 20-week-old male *Lpcat1*-WT and *Lpcat1*-KO mice (includes data also plotted in Figures 4A-4C).

(C-E) PC and PE compositions of control (C), injured (barium chloride injection, 14 dpi) (D), or dystrophic (*mdx* background) (E) TA muscles of 20-week-old male *Mboat2*-WT and *Mboat2*-KO mice (includes data also plotted in Figures 4D-4I).

Significance is based on t-tests; n = 4–10 / group.

**Figure S5. Gene expressions and PC and PE compositions of neonatal *Mboat2*-WT and -KO muscles, related to Figure 6**.

Gene expressions and PC and PE compositions were measured in gastrocnemius muscles of male adult (1-year-old) and neonatal (1-week-old) *Mboat2*-WT and *Mboat2*-KO mice. All mice were born in mixed litters from *Mboat2*-heterozygous mothers, to avoid maternal influences on the analyses of neonatal tissues.

(A-D) Gene expressions were measured for *Mboat2* (A) and satellite cell markers *Pax7* (B), *Myod1* (C), and *Myogenin* (D) (n = 4-7 / group).

(E) Body weights of *Mboat2*-WT and -KO male mice (n = 5-13 / group).

(F-G) PC and PE compositions of gastrocnemius muscles of 1-year-old (F) or 1-week-old (G) *Mboat2*-WT and -KO mice (n = 7-8 / group; includes same PC data plotted in figures 6A-6C).

Error bars represent SD (A-G). Significance is based on t-tests (F,G).

**Figure S6. Dose response of AAV-*Mboat2* to increase PC 34:1 in muscle and supportive evidence for membrane pacemaker theories, related to Figure 7.**

AAV-*eGFP* particles were injected into right gastrocnemius muscles and AAV-*Mboat2* expression particles were injected into left gastrocnemius muscles of 5- to 8-week-old female WT-B10 mice. Each expression particle was injected at 4 different doses (n = 1 muscle / dose). Tissues were collected 15 days after injection for analyses by LCMS and RT-PCR.

(A) PC compositions of muscles injected with AAV-*Mboat2* particles.

(B) PC compositions of muscles injected with AAV-*eGFP*. (C-D) Expressions of AAV-delivered transgenes, *Mboat2* (C) and *eGFP* (B), in the injected tissues.

Table S1. Fourteen mammalian LPLATs

| LPLAT name [3] | gene name | |
|---|---|---|
| | mouse | human |
| LPLAT1 | *Agpat1* | *AGPAT1* |
| LPLAT2 | *Agpat2* | *AGPAT2* |
| LPLAT3 | *Agpat3* | *AGPAT3* |
| LPLAT4 | *Agpat4* | *AGPAT4* |
| LPLAT5 | *Agpat5* | *AGPAT5* |
| LPLAT6 | *Lclat1* | *LCLAT1* |
| LPLAT7 | *Lpgat1* | *LPGAT1* |
| LPLAT8 | *Lpcat1* | *LPCAT1* |
| LPLAT9 | *Lpcat2* | *LPCAT2* |
| LPLAT10 | *Lpcat4* | *LPCAT4* |
| LPLAT11 | *Mboat7* | *MBOAT7* |
| LPLAT12 | *Lpcat3* | *LPCAT3* |
| LPLAT13 | *Mboat2* | *MBOAT2* |
| LPLAT14 | *Mboat1* | *MBOAT1* |

Table S2. Phosphatidylcholine (PC) alterations reported in dystrophic muscle

| Subjects | Changes in dystrophic muscle | Reference |
|---|---|---|
| DMD patients | ↑ 18:1, ↓ 18:2 in PC | [4, 5] |
| DMD patients | ↑ 18:1 in PC | [6] |
| DMD patients | ↑ PC 34:1 / PC 34:2 ratio | [7] |
| DMD patients | ↑ PC 34:1 | [8] |
| *mdx* mice | ↑ PC 34:1 / PC 34:2 ratio | [9, 10] |
| *mdx* mice | ↑ 18:1 in total phospholipids | [11] |
| *mdx* mice | ↑ PC 34:1 | [12] |
| *mdx* mice | ↑ PC 34:1, ↑ PC 34:2 | [13] |
| *mdx* mice | ↑ PC 34:1, PC 34:2 variable | [14] |

Table S3. Estimated fatty acids in diets (% of total fatty acids by weight)

| fatty acid | high 18:1 diet | high 18:2 diet | regular chow |
|---|---|---|---|
| 16:0 | 12.8 | 13.2 | 18.2 |
| 16:1 | 0.1 | 0.1 | 1.0 |
| 18:0 | 3.8 | 3.2 | 2.5 |
| 18:1 | 55.0 | 19.7 | 22.3 |
| 18:2 | 24.2 | 59.6 | 46.0 |
| 18:3 | 3.0 | 3.1 | 3.4 |
| 20:5 | 0.0 | 0.0 | 2.4 |
| 22:6 | 0.0 | 0.0 | 2.2 |
| Saturated | 17.6 | 17.4 | 21.6 |
| Monounsaturated | 55.1 | 19.9 | 23.8 |
| Polyunsaturad | 27.2 | 62.7 | 54.6 |
| n-3 | 3.0 | 3.1 | 8.5 |
| n-6 | 24.2 | 59.2 | 46.2 |

Fatty acids comprising ≥ 0.5% are shown. Custom diet values reported by manufacturer. Standard chow values measured by GC-FID.

Table S4. Nutrition of diets

| | high 18:1 diet | high 18:2 diet | regular chow |
|---|---|---|---|
| kcal% protein | 20.1 | 20.1 | 30.1 |
| kcal% fats | 15.8 | 15.8 | 12.2 |
| kcal% carbs | 64.2 | 64.2 | 57.7 |
| kcal / 100 g | 399 | 399 | 339 |

Nutritional components were reported by manufacturers.
Energy contents calculated with Atwater formula.

| Table S5. Enzymatic activities of selected LPLATs | | | | |
|---|---|---|---|---|
| LPLAT | preferred substrates | | main regulated products | references |
| | acyl-CoA | lysophospholipid | | |
| LPCAT1 | 16:0-CoA | LPC | saturated PC | [15-19] |
| LPCAT3 | 20:4-CoA | LPC, LPE | PUFA-containing PC and PE | [20] |
| MBOAT2 | 18:1-CoA | LPC, LPE | MUFA-containing PC and PE | [21] |
| AGPAT3 | 22:6-CoA | LPA | DHA-containing-PC and -PE | [22-24] |

| Table S6. List of primers used for RT-PCR | | |
|---|---|---|
| Gene | Forward primer | Reverse primer |
| *Agpat1* | AAACGAGGCGCCTTCCA | GGAGTAGAAGTCTTGATAGGAGGACATG |
| *Agpat2* | TGTGGGCCTCATCATGTACCT | AGGTCGGCCATCACAGACA |
| *Agpat3* | AAGCACCTATACCGCCGTATCA | GACCACCACTCCAGGAGCAT |
| *Agpat4* | AAGCAGCTGTTCCGCAAGA | CCACCACTCCAGAAGCATCA |
| *Agpat5* | AATGAGAAAGGTTCAGGAAAATACTCA | TGAATATGAAGTTTTGGGCACTGT |
| *Lclat1* | TGGATGTTCCTGTGGAACTGTCT | GGTTCATGGATGGCACAAAAATA |
| *Lpgat1* | TCCCAAAGCTGAACCAATAGACA | AATAAAGCGCTGATAGAGCCAGC |
| *Lpcat1* | TCCCAGACCTTAGCCACCAT | ACAGGTTGGCCTCATCTATGCT |
| *Lpcat2* | TCCACGTTCTTCGACGGAAT | TGGCTGCAAAGCCCGTAA |
| *Lpcat4* | AGAGGGTTAAGTTCTGCCTCCT | CATACAGTCTTCCTCCATCCTGTAA |
| *Mboat7* | ATACTGGAACATGACCGTGCAGT | TAGGTAGTAACCAGGGTGGAGGC |
| *Lpcat3* | AGATGGAATTCCTCATTGTTATCG | GAAGGGCTGTAGGGCAGTGA |
| *Mboat2* | CTGCAGCCCCTCAGCAA | CGGCTGCTAACAAGGCAAAG |
| *Mboat1* | CTGAAATGTGTGTGCTATGAGCG | TGGAAGAGAGGAAGTGGTGTCTG |
| *Pax7* | TCAAGCCAGGAGACAGCTTG | TGTGGACAGGCTCACGTTTT |
| *Myod1* | AGCATAGTGGAGCGCATCTC | GTTCCCTGTTCTGTGTCGCT |
| *Myogenin* | GTGAATGCAACTCCCACAGC | CCACGATGGACGTAAGGGAG |
| *Myf6* | GTGGACCCCTACAGCTACAAA | ACGATGGAAGAAAGGCGCT |
| *18S* | CTCAACACGGGAAACCTCAC | AGACAAATCGCTCCACCAAC |
| *eGFP* | GCATCGACTTCAAGGAGGAC | GAAGTTCACCTTGATGCCGT |
